# Glucose controls co-translation of structurally related mRNAs *via* the mTOR and eIF2 pathways in human pancreatic beta cells

**DOI:** 10.1101/2021.09.13.460006

**Authors:** Manuel Bulfoni, Costas Bouyioukos, Albatoul Zakaria, Fabienne Nigon, Roberta Rapone, Laurence Del Maestro, Slimane Ait-Si-Ali, Raphaël Scharfmann, Bertrand Cosson

**Affiliations:** Université de Paris, CNRS UMR7216 Epigenetics and Cell Fate, Paris F-75013, France; U1016 Inserm, Institut Cochin, 75014 Paris; Institut de Biologie de l’École Normale Supérieure ENS. CNRS UMR8197. Inserm U1024. 75005 Paris, France

## Abstract

Pancreatic beta cell response to glucose is critical for the maintenance of normoglycemia. A strong transcriptional response was classically described in rodent models but, interestingly, not in human cells. In this study, we exposed human pancreatic beta cells to an increased concentration of glucose and analysed at a global level the mRNAs steady state levels and their translationalability. Polysome profiling analysis showed an early acute increase in protein synthesis and a specific translation regulation of more than 400 mRNAs, independently of their transcriptional regulation. We clustered the co-regulated mRNAs according to their behaviour in translation in response to glucose and discovered common structural and sequence mRNA features. Among them mTOR- and eIF2-sensitive elements have a predominant role to increase mostly the translation of mRNAs encoding for proteins of the translational machinery. Furthermore, we show that mTOR and eIF2α pathways are independently regulated in response to glucose, participating to a translational reshaping to adapt beta cell metabolism. The early acute increase in the translation machinery components prepare the beta cell for further protein demand due to glucose-mediated metabolism changes.

**AUTHOR SUMMARY:** Adaptation and response to glucose of pancreatic beta cells is critical for the maintenance of normoglycemia. Its deregulation is associated to Diabetic Mellitus (DM), a significant public health concern worldwide with an increased incidence of morbidity and mortality. Despite extensive research in rodent models, gene expression regulation in response to glucose remains largely unexplored in human cells. In our work, we have tackled this question by exposing human EndoC-BH1 cells to high glucose concentration. Using polysome profiling, the gold standard technique to analyse cellular translation activity, we observed a global protein synthesis increase, independent from transcription activity. Among the specific differentially translated mRNAs, we found transcripts coding for ribosomal proteins, allowing the cell machinery to be engaged in a metabolic response to glucose. Therefore, the regulation in response to glucose occurs mainly at the translational level in human cells, and not at the transcriptional level as described in the classically used rodent models.

Furthermore, by comparing the features of the differentially translated mRNAs, and classifying them according to their translational response, we show that the early response to glucose occurs through the coupling of mRNA structure and sequence features impacting translation and regulation of specific signalling pathways. Collectively, our results support a new paradigm of gene expression regulation on the translation level in human beta cells.

## INTRODUCTION

Pancreatic islet β-cells play a pivotal role in the maintenance of normoglycemia by synthesizing, storing and secreting insulin. Glucose uptake and metabolism are essential for regulation of glycemia by stimulating insulin secretion and triggering specific gene expression. These processes have been widely studied in rodent since 1970s (1). Due to the long-standing difficulties to generate a human cellular model (2), or to access to primary human islet preparations derived from deceased donors (3), there is a scarcity of results obtained from human cells. In addition, despite many similarities, there are major differences between human and rodent models such as the copy number of *insulin* genes, the expression of different transcription factors in glucose-stimulated insulin secretion, the architecture of the Islet of Langerhans with functional implication and, finally, the susceptibility to β-cell injury (2). Moreover, concerning glucose-dependent gene expression regulation, recent transcriptome studies have demonstrated important differences in expression levels between human and rodent cell lines. In the rat β-cell line INS-1, more than 3700 genes were significantly affected in response to glucose (4). In contrast, a recent transcriptome study of the first human β-cell line (EndoC-BH1), able to secrete insulin in response to glucose stimulation (5), showed that only a scarce number of genes were modified at the mRNA level for cells exposed for eight hours either to high or low glucose concentrations (6). Accordingly, a previous report on donor human islet treated similarly during twenty-four hours (7) reported that the expression of only 20 genes was affected. Taken together, these findings highlight a considerable difference in gene expression regulation at transcriptional level between human and mouse pancreatic β-cell.

Glucose regulation has also been addressed at post-transcriptional level in rodent models since the 70s. In particular, attention was focused on the regulation of glucose-induced pro-insulin synthesis (8–10) reporting that the first cellular response to replenish insulin was entirely mediated at translational level without affecting mRNA abundance (8). Beside *pro-insulin*, synthesis of other proteins was also stimulated, but these proteins were not identified. A translatome study addressed this issue in mouse insulinoma 6 (MIN6) cell line by polysome profiling (11) and identified 313 mRNAs, for which the association with polysomes was changed by at least 1.5 times. Interestingly, in low glucose the Integrated Stress Response (ISR) mediates eIF2 phosphorylation, promoting translation of a group of mRNAs including b-zip transcription factors such as ATF4, CHOP (DDIT3), and c-Jun. The translation of these mRNAs was reduced upon glucose increase and eIF2 dephosphorylation, linking regulatory pathway activity and protein expression regulation (11).

To date, no translatome studies have been made on human β-cells to address glucose induced post-transcriptional regulation. In this work, we exposed human pancreatic β-cells exhibiting glucose-inducible insulin secretion (12) to high glucose concentrations for 30 min to observe the early cellular response. We observed a global protein synthesis increase, independent from transcription regulation. We identified 402 mRNAs that are differentially translated in response to glucose and identified different groups of co-regulated transcripts. We found mTOR and eIF2-sensitive elements in a majority of them and, accordingly, we found that both pathways activate translation of specific mRNAs in response to glucose. Upregulated genes are mainly coding for ribosomal proteins, increasing the translation machinery potential, allowing the cell machinery to be engaged in a metabolic response to glucose.

## RESULTS

### Glucose induces an acute increase in protein synthesis in human β-cells

To monitor and quantify the translation activity in response to glucose, we performed a polysome profiling analysis on a functional human β-cell line able to produce insulin in response to glucose (EndoC-βH2 cells, see Material and Methods). Polysome profiling is a gold standard technique to analyze cellular translation activity. Ribosome complexes of different densities are separated on a sucrose gradient consistent with the number of ribosomes. Briefly, EndoC-βH2 cells were cultured for 24 hours in 0.5 mM glucose and then treated for 30 min with either 0.5 mM or 20 mM glucose. Polysome profiling showed that increasing glucose to 20 mM triggered an important increase in the content of polysomes in parallel to a decrease of the monosomes peak (80S) (Fig. 1A). Polysome/monosome ratio was raised 2 times, which is classically observed with a global increase in protein translation. This translation upregulation was confirmed by ^35^S-Met incorporation (Supp. Fig 1A).

**Figure 1.**
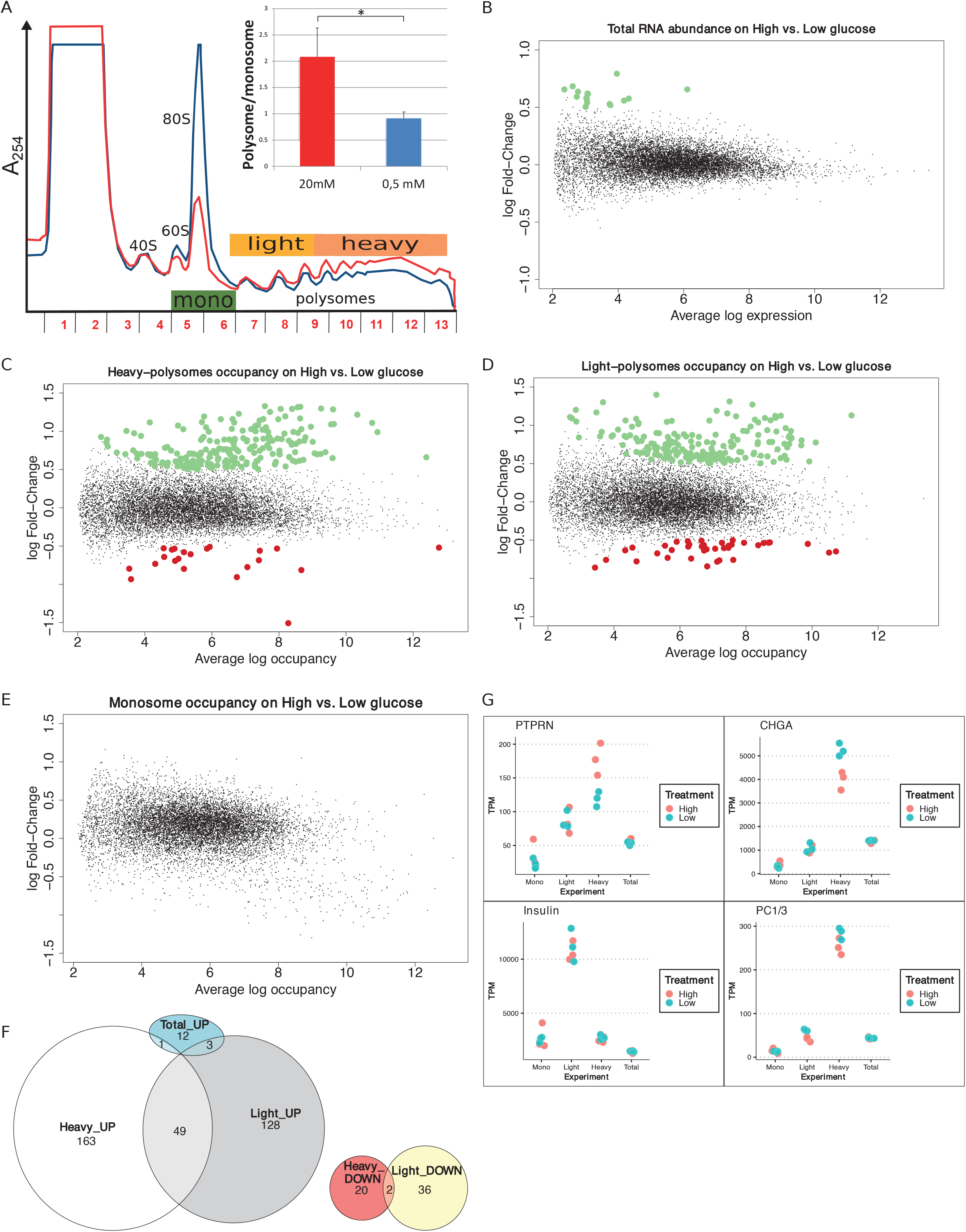
Glucose induces increase in protein synthesis without affecting mRNA abundance and regulates translation rates of a subset of mRNAs. **A**. Polysome profiles of EndoC-βH2 with Low glucose (blue) and High glucose (red). The absorbance at 254 nm (A254) recorded during the collection of the fractions of the gradient is displayed. The positions of 40S, 60S, 80S and polysomes are indicated. The tree-colored bars represent the fractions that were pooled for sequencing: monosomes (green), light polysomes (yellow), heavy polysomes (light orange). EndoC-βH2 cells in high glucose had a significantly higher polysome/monosome ratio than did EndoC-βH2 cells in low glucose. Cells were treated in parallel using paired culture plates, and centrifugated together in the same rotor. Figure shows a representative replicate. The statistical significance of the polysome/monosome ratio was assessed using a paired t-test from three independent experiments (*, p < 0.001, n = 3). **B**. MA-plot of total mRNA abundance, dots in green specify up-regulation. **C-E**. MA-plots for each pool of fractions: heavy polysomes (C), light polysomes (D) and monosomes (E); dots in green corresponds to transcripts that are upregulated upon glucose stimulation, dots in red to transcripts that are downregulated. **F**. Venn diagram showing the overlap between glucose UP- or DOWN-regulated transcripts in Light and Heavy polysomes. Total_UP are transcripts varying in abundance. **G**. TPM values for each condition and each replicate are plotted. In red High glucose replicates, while in cyan Low glucose replicates. Names of the plotted gene is indicated above.

We next quantified whether the global increase in translation was associated to a modification in mRNA steady state levels by genome-wide transcriptome analysis. As illustrated by the MA plot in Fig. 1B, global mRNA levels were not significantly affected by a 30 min glucose treatment. Only 16 transcripts (Supp. Fig 1B) were found to be significantly affected, 7 of them coding for histone proteins. However, these variations are low, with a fold change of less than 2-fold. We validated this result by RT-qPCR showing that *Histone1H3C* and *Histone1H3D* mRNAs increased from 30 min after glucose shift (Supp. Fig.1C), in an extent similar to that observed by RNAseq (log FC = 0.7 at 30 min, Supp. Fig 1B, corresponding to FC = 1.6). Conversely, the abundance of mRNAs such as, *PTPRN, CHGA* or *CCNG1* was not significantly affected even after 1h or 2h after the glucose shift. This finding is in agreement with the results described in Richards et al., showing that even 8 hours after glucose increase, only a scarce number of genes were modified at the mRNA level (6). Hence, the global increase in protein synthesis is virtually independent from transcription regulation.

In conclusion, we show here for the first time that human beta cells respond to glucose by a rapid and important increase in mRNA translation, which is virtually independent of changes in the transcriptome.

### Glucose regulates translation rates of mRNAs involved in the insulin secretion pathway and in the translation machinery

We next studied genes whose translation rates are regulated in response to glucose by sequencing the mRNAs associated with monosomes and polysomes (Fig. 1A, see Materials and Methods). The commonly adopted strategy to identify translated genes is usually to consider mRNAs associated with more than 3 ribosomes (13), but each fraction from monosomes to heavy polysomes could also be sequenced (14). To obtain a good compromise between resolution and sensitivity, we analyzed separately the monosomes (80S), light polysomes (2-4 ribosomes per mRNA), and heavy polysomes (> 4 ribosomes per mRNA). Accordingly, fractions were pooled (see Fig. 1A) to collect monosomes (fractions 5-6), light polysomes (fractions 7-9) and heavy polysomes (fractions 10-13) before RNA sequencing.

Differential analysis using the limma package (15) highlighted that the abundance of 235 mRNAs in heavy polysomes and 218 in light polysomes was significantly changed upon glucose shift (adjusted p-value < 0.05 and logFC > 0.5, Fig 1C and 1D), but not in monosomes (Fig. 1E). These changes do not correlate with a significant difference in mRNA total level (Fig. 1B). As expected from the global increase in translation we observed in high glucose (Fig. 1A), most of the identified transcripts were enriched in polysomes (Fig. 1F): 164 in the heavy polysomes, 131 in the light polysomes, 49 common to both. Thus, only a minority of transcripts, 58 in total, were downregulated. Interestingly, from the 16 transcripts increased in RNA abundance (Fig 1B) only four have been found increased in polysomal fractions (Fig. 1F and Supp. Fig. 1A). Except for these four genes, the variation for each transcript observed in polysomal fractions corresponds to a specific translational regulation independently of any variation in mRNA abundance and, consequently, of any transcriptional regulation.

Continuing the investigation of differentially translated genes, we proceed by assessing if glucose stimulation modified the biosynthesis of insulin and of known major factors involved in insulin maturation and secretion pathway. We focused on gene products whose biosynthesis was reported to be enhanced in response to glucose in rodent cells, such as Protein Tyrosine Phosphatase Receptor Type N (PTPRN) (16), chromogranin A (CHGA) (17), and pro-hormone convertases 1/3 (PC1/3) (18). In heavy polysomes, *PTPRN* mRNA was enriched while *CHGA* mRNA was reduced (Fig. 1G and Supp. Fig. 2C and 2D). Interestingly, the relatively short *Insulin* transcript associates mainly with light polysomes (Fig. 1G), as observed previously in a mouse cellular model (11).

Our data show that the human EndoC-βH2 beta cell line incubated with high glucose for 30 min quickly modify the translation rates of at least two mRNAs that code for proteins involved in the insulin secretion pathway. In contrast, We did not find any particular expression difference in the dot plot for *Insulin* and *PC1/3* mRNAs (Fig. 1G), and accordingly we did not find these genes as differentially translated in our conditions.

Gene Ontology (GO), gene set and REACTOME pathway enrichment analyses were performed for all the Differentially Translated Genes (DTGs). Figure 2 illustrate the gene-concept plots (cnetplots) of each ontology term and all its associated genes of the top 10 enriched categories for molecular functions (MF, Fig. 2A) and biological processes (BP, Fig. 2B). Cnetplots are an informative way to represent the relationships between different ontology terms in a graph of all the associated genes together with the logFC for each gene. It is evident that the majority of the translationally upregulated genes (red nodes) are enriched in GO terms related to the biosynthesis and metabolism of proteins, rRNAs and to the translation machinery (Fig. 2). Supp. Fig. 2E illustrates the top 20 REACTOME pathways that are enriched in DTGs which further corroborate the finding of the GO analysis (Fig. 2).

**Figure 2.**
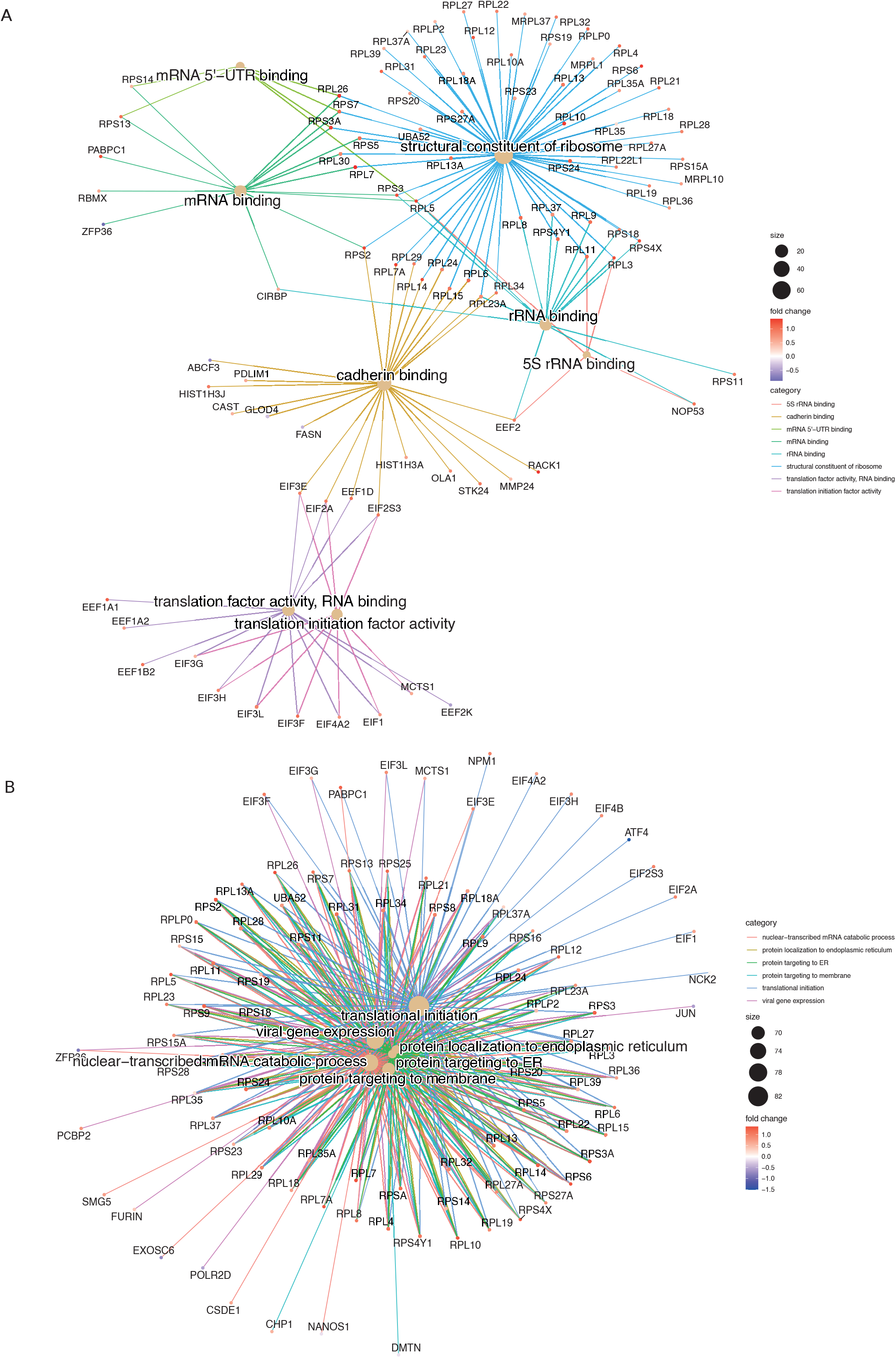
Gene Ontology enrichment analysis, gene-concept plots (cneplots). Each cneplot illustrates the most significantly enriched Molecular Functions (MF, **A**) and Biological Processes (BP, **B**) categories. Lines of the same colour connect genes of the same GO category and the colour gradient of each gene corresponds to its log-FC.

In conclusion, our results show that glucose concentration changes modulate translation rate of specific mRNAs of the insulin secretion pathway and promotes the synthesis of the translation machinery components.

### Differentially translated mRNAs in response to glucose share unique structural and sequence features

It might be tempting to explain the translation modulation of specific mRNAs in response to glucose by a simple global translation increase. To test this, we calculated the translation ratio, defined as the abundance ratio of the translating mRNA in mono/polysome fractions to total mRNA regarding a certain gene (see Material and Methods for detail), for the most translated transcripts for each gene (Supp. Table 1), and then asked if the 200 most translated mRNAs are the same in low and high glucose (Fig. 3A), by ranking them according to the translation ratio. We found that around 40% (76) of the best translated transcripts in high glucose (Fig. 3A, grey circle) were not among the best translated in low glucose (Fig. 3A, blue circle). Furthermore, 80% of the mRNAs that display the major differences in ranking (Fig. 3A, white circle) are not found in the 200 top translated mRNAs in high or low glucose. The cellular response to glucose is therefore not just a simple increase in overall translation.

**Figure 3.**
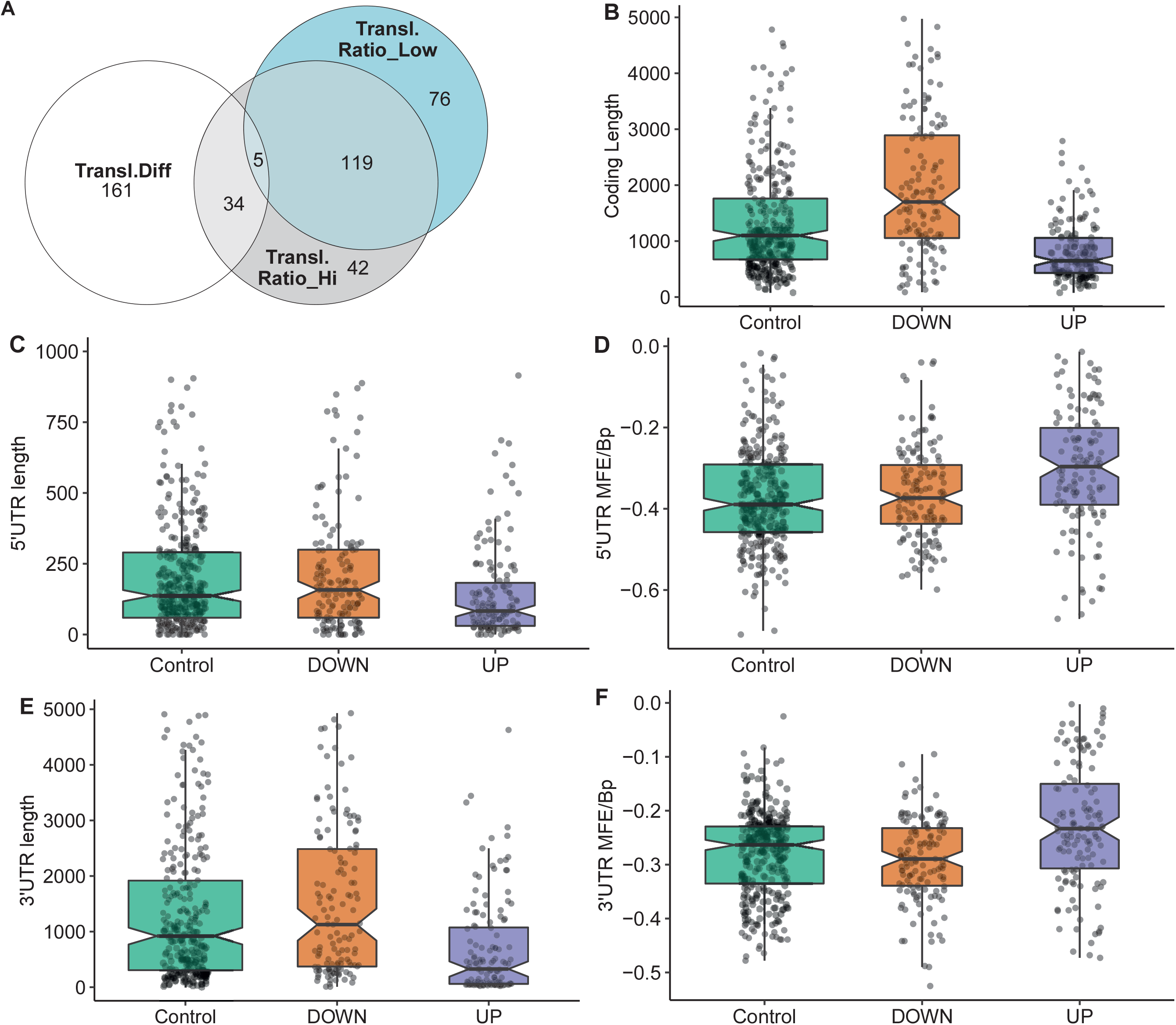
mRNA feature analysis of the 3 groups of differentially translated mRNAs. **A**. Venn diagram showing the overlap between the 200 mostly translated genes: best translated in high glucose (grey circle), best translated in low glucose (blue circle), mRNAs with the most difference ones in ranking between low and high glucose (white circle). **B-F**. mRNA features analyses of 3 groups of mRNAs based on the difference in translation ratio between high and low glucose. Higher (UP) or the lower (DOWN) translation ratio in high glucose, and a control group with no significant changes between high and low glucose. MFE/Bp is the folding free energy normalized by the length (see Material and Methods). **B**, coding length; **C**, 5’UTR length, **D**, 5’UTR MFE/Bp, **E**, 3’UTR length, **F**, 3’ UTR MFE/Bp.

Translational adaptation to glucose seems therefore to be a complex process. Thus, we further investigated potential association between the differential translation regulation and specific sequence and structural features of the DTGs. To this end, we have classified differences in translation ratios between high and low glucose to 3 mRNA groups: 147 mRNAs with higher translation ratio in high glucose, 137 with a lower translation ratio in high glucose and a control category of 320 mRNAs with no significant change between high and low glucose. The most translated transcript of each of these 604 genes was analysed by our in-house developed software (detailed in Materials and Methods) to retrieve sequence information from ENSEMBL database and identify structural features. Several characteristics appear in the statistical analysis of the features (Fig. 3B-F and Supp. Fig. 3). Among them, the minimum folding energy normalized over the length of the sequence (MFE per BP) is an interesting measure to estimate the complexity of an mRNA untranslated region (UTR) structure.

Transcripts with higher translation ratio difference between high and low glucose appear to have statistically significant shorter open reading frames (ORFs), and shorter and less complex UTRs. Conversely, transcripts that are down regulated have longer ORFs and more structured 3’UTR since their MFE per BP distribution slightly decreases (Fig. 3 B-F).

These results indicate that the sequence and structural features we have used allow the classification and characterization of the highly translated mRNAs; so they provide a good tool to dissect the different classes of translational behaviour of transcripts in mono-, light- and heavy-polysomes. Motivated by that, we proceeded with the dissection of this behaviour and the sequence-structure characterization of the different transcript classes.

### Differentially translated mRNA clusters display specific mRNA features

To group the translationally co-regulated transcripts, we clustered the 402 differentially translated mRNAs we identified previously (Fig. 1F) based on their behaviour between monosomes, light, and heavy polysomes. To this end, we calculated for each mRNA the log2 ratio of the average abundance in high over low glucose condition for each of the three ribosomal fractions. The log ratios matrix was then subjected to a modelling clustering method able to determine in an unsupervised way the best model that characterizes the data (19). The model generated six clusters, which highlighted six different types of behaviours (Fig. 4A, coloured bars with the cluster number). Genes of the top cluster (n°1, seagreen bar) showed a clear pattern for 73 mRNAs shifting from the monosomes to polysomes (both light and heavy), which is the expected behaviour for a mRNA with increased translation. Cluster n°6 (102 mRNAs) presented a behaviour similar to cluster 1 with an increase of mRNAs in heavy polysomes but without any rise in the light polysomes. These transcripts move from the monosomes to the heavy polysomes fraction and correspond to transcripts whose translation is greatly increased. Instead, the mRNAs of clusters 2 and 3 (90 and 79 mRNAs, respectively) showed increased mRNA levels for light/heavy and monosome/light polysomes, respectively. We reasoned that at low glucose concentration these mRNAs are associated with small complexes that have a density smaller than one ribosome and correspond to mRNAs newly recruited to the translation machinery. Finally, clusters 4 and 5 (37 and 21 mRNAs, respectively) collected all the mRNAs whose levels in the polysome fractions decreased upon glucose stimulation, which might reflect a decrease in their translation. The difference between these two clusters comes from the fact that the decrease is observed in light polysomes for cluster 4 and in heavy polysomes for cluster 5.

**Figure 4.**
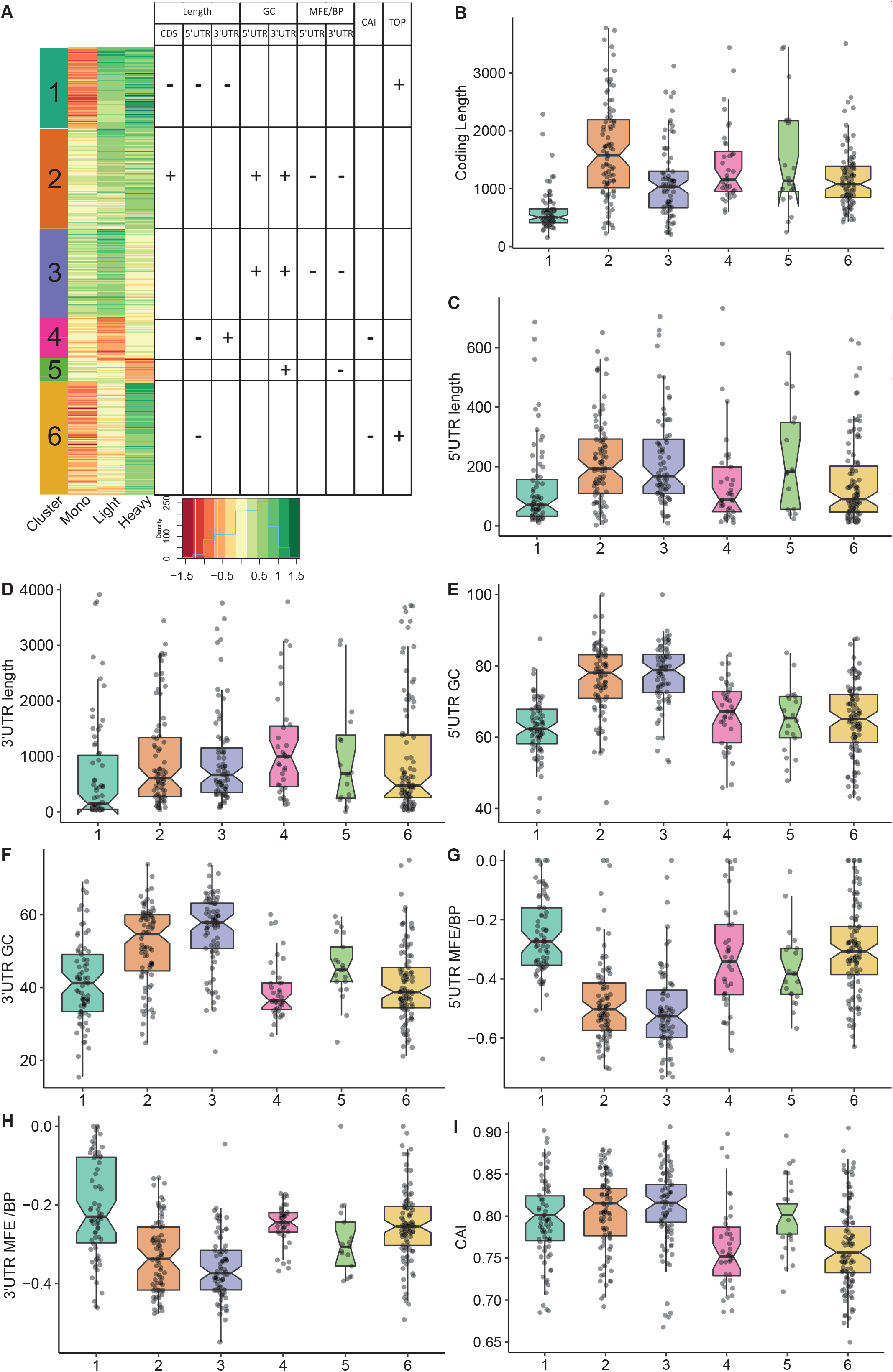
Clustering and mRNA features analysis. **A**. Heatmap of the log2 ratios between the average TPM of each differentially translated gene in high-over low-glucose. An unsupervised model clustering algorithm was used to cluster the differentially translated genes into six groups represented here with the coloured vertical bar at the left side of the figure. Colours range from dark red when genes are less represented in a polysome fraction to dark green when overrepresented in a polysome fraction in high glucose condition. The table summarizes the variations observed for the different features presented in **B-I**: Box plots of the different mRNA features of the six-identified cluster.

Based on the link between features and behaviours that we observed (Fig. 3), we hypothesized that the different patterns observed for the clusters could be a consequence of *cis*-regulatory elements present on the mRNAs which could serve as binding sites for *trans-*acting factors such as RNABPs and miRNAs. To investigate this possibility, we calculated a series of mRNA features for the most abundant transcript for each gene. We developed an RNA feature extraction tool (see Material and Methods) that is able to download, from the ENSEMBL database bioMart (20), the transcript sequences with additional annotations and calculate sequence and structural properties (see Supp. Table 2 for the full table of results). We proceeded by grouping these mRNA features between the 6 clusters of different translation behaviour identified by the model clustering approach (Fig. 4 B-I, the full ensemble of the statistical analysis of differences between groups, including the results of all the Kruskal-Wallis H-test and its associated p values are available in supplementary Supp. Fig. 4). Strikingly, the length of the coding sequence (CDS) of cluster 1 mRNAs was significantly shorter, while in cluster 2 the CDS were longer (Fig. 4B). Next, we analysed the length of the UTRs (Fig. 4C-D). Notably, there was a tendency for mRNAs of cluster 1 to have shorter 5’ and 3’ UTRs than the other clusters. Cluster 4 and 6 showed similar tendencies towards shorter 5’ UTRs (Fig. 4C). Cluster 4, that corresponds to mRNAs whose translation decreases in response to glucose, contained longer 3’ UTRs (Fig. 4D).

We next analysed the GC-richness of the 5’ and 3’UTRs, as a proxy to determine the complexity of secondary structures formed by the mRNA UTRs. Cluster 2 and 3 contained mRNAs with higher GC content on both 5’ and 3’ UTRs than the others (Fig. 4E-F). In accordance, clusters 2 and 3 contained mRNAs with more structured 5’ and 3’ UTRs than the other clusters (Fig 4G-H). In addition, cluster 5, the cluster with the underrepresented mRNAs in the heavy polysome fraction, also appears to have the next higher GC content and lowest MFE per BP on its 3’UTR (Fig. 4F,H), indicating that the 5’UTR structural and sequence complexity affects the translation regulation.

In addition to the UTR structure, we found a tendency towards a lower Codon Adaptation Index (CAI) for cluster 4 and cluster 6 (Fig. 4I). Since CAI is associated with slowest translation rates this finding could explain at least in part the accumulation of transcripts in monosomes (cluster 6) or light polysomes (cluster 4) rather than in heavy polysomes in low glucose (Fig. 4A).

We then searched for functional motifs in the UTRs by interrogating the UTRdb (http://utrdb.ba.itb.cnr.it/,(21), and we found already characterized RNA binding motifs in the UTRs of the mRNAs which are differentially translated (Supp. Fig. 5). Interestingly, we found TOP motifs (5’- terminal oligopyrimidine (TOP) (22)) mostly represented in cluster 1 and 6 (Fig. 5A), and also uORFs, and IRES features (Fig. 5B). It is interesting to note that uORF and TOP features are mutually exclusive (Fig.5B), meaning that these features are used independently to differentially regulate mRNA translation. This finding indicates two different potential modes of translation regulation from two independent mechanisms.

**Figure 5.**
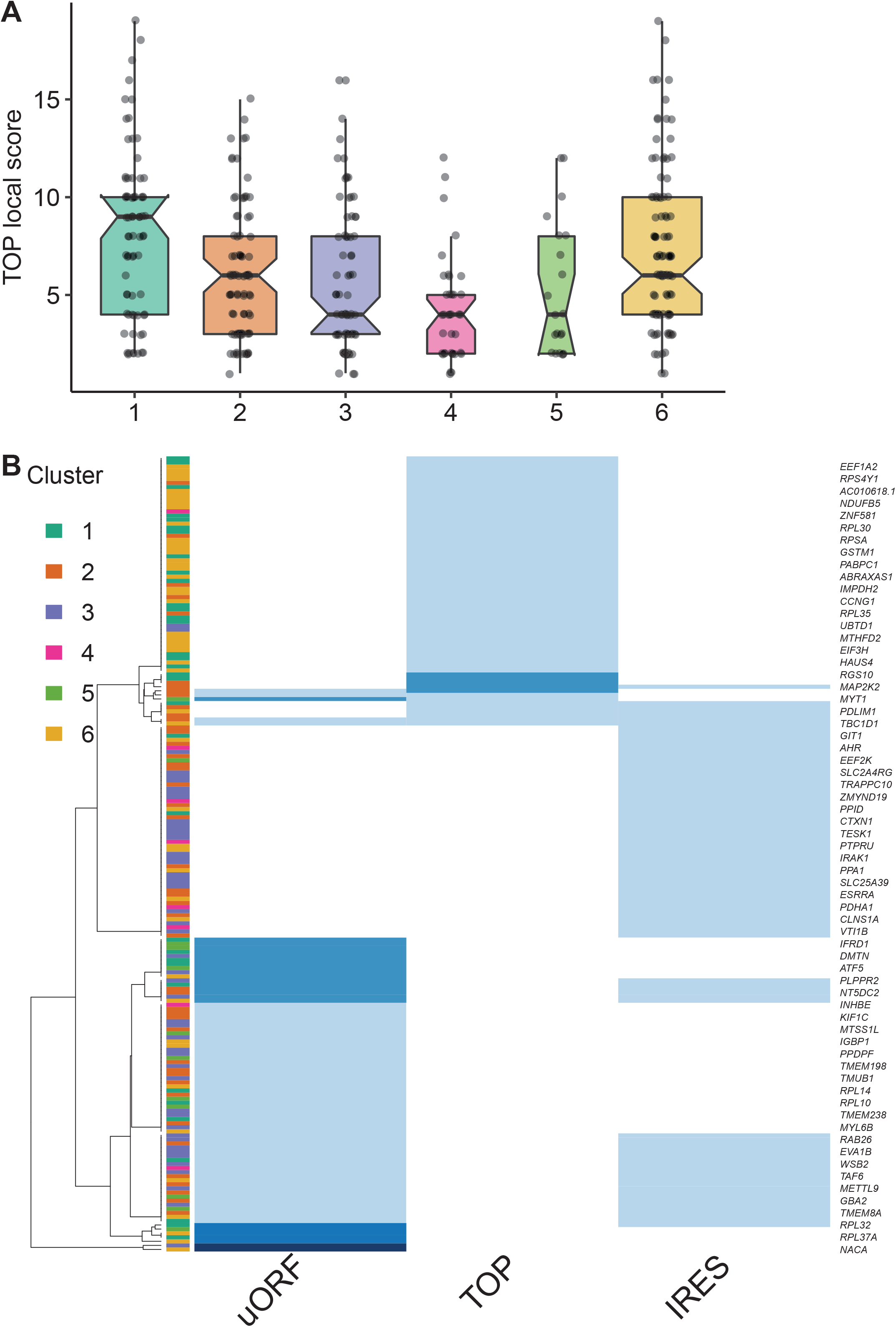
Functional motifs analysis of the UTRs by interrogating the UTRdb for TOP motifs (A), and TOP, uORF and IRES in the 5’UTRs (B), common gene names of the encoding gene of each transcript are indicated on the right side. The darker the blue, the higher the number of features per RNA (from 1 to 3 per RNA).

Taken together, the results suggest that the translational regulation of differentially translated mRNAs upon glucose concentration shift is strongly associated with, and can be characterized by, specific sequences as well as structural features on the UTRs and coding regions of these mRNAs.

### Clustering reveals that specific pathway regulators are co-regulated in response to glucose, and that TOP motifs are preponderant to upregulate the translation machinery components

Next, we performed gene ontology enrichment analyses, using the R package ClusterProfiler (23), in order to assess if the mRNAs belonging to the different clusters were implicated in specific biological processes. Cluster 1 and 6 were enriched for categories related to translation but especially for ER-related translation (Fig. 6A-B). Cluster 4 (Fig. 6C), which contained the translationally repressed mRNAs in high glucose, was enriched in mRNAs coding for proteins involved in regulation of metabolic processes and cell death. In particular, genes involved in tricarboxylic acid, acyl-CoA, and thioester metabolisms were found to be enriched in cluster 4. Cluster 5 with also repressed transcripts in high glucose was enriched for categories related to response to starvation and growth factors (Fig. 6D), including mRNAs coding for b-zip transcription factors such as ATF4, DDIT3(CHOP), and c-Jun (Supp. Table 2) that had a similar behaviour in mouse beta cell (see introduction,(11)).

**Figure 6.**
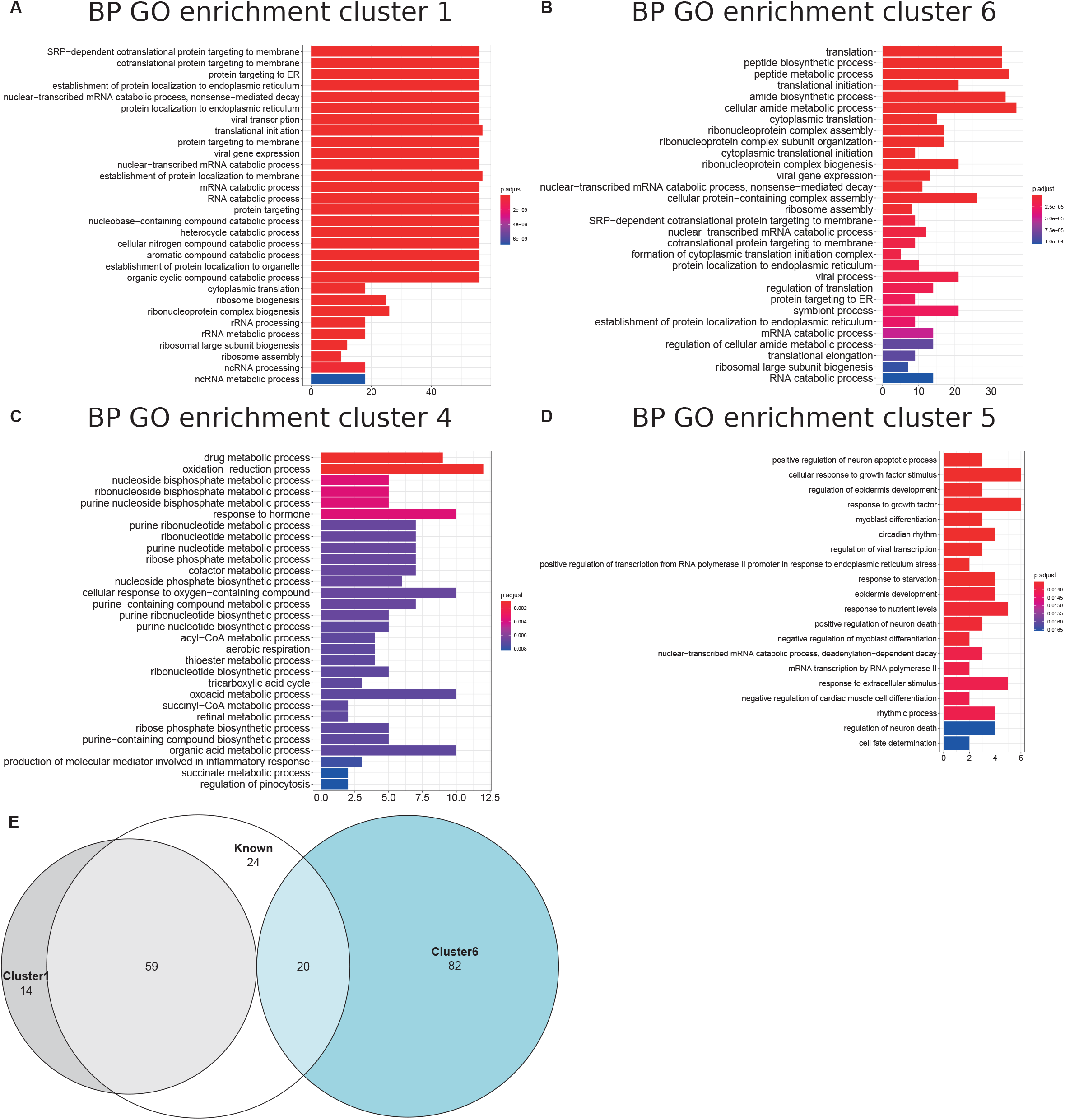
Cluster Gene Ontology enrichment. **A-D**. barplot of gene ontology terms enriched for biological processes for clusters 1, 4, 5 and 6 using the R package clusterProfiler. **E**. Venn diagram of genes belong to cluster 1 (grey circle) and 6 (blue circle) with previously reported TOP-RNAs (white circle).

We thus investigated if ribosomal proteins or translation factors were found in clusters 1 and 6. We found that cluster 1 contained 56 mRNAs coding for ribosomal proteins (Supp. Table 3, RPL and RPS), corresponding to 70% of the total number of RP transcripts, while cluster 6 contained 13 mRNAs coding for eukaryotic translation factors of which 4 were elongation factors (eEF1A1, eEF1A2, eEF1B2, eEF2) and the remaining were initiation factors (eIF2A, eIF2S3, eIF3E, eIF3F, eIF3G, eIF3H, eIF3L, eIF4A2, eIF4B) (Supp. Table 3, eIFs and eEFs). These mRNAs are known to contain the TOP motif (TOP-RNAs, (22,24,25)), that we found enriched in the differentially regulated mRNAs (Fig. 5A), mostly in cluster 1 and at a lesser extent in cluster 6. Therefore, we compared the mRNAs found in cluster 1 and 6 with the list of mRNAs that were previously reported or proposed to be TOP-RNAs in three already published studies (22,24,25) (Supp. Table 4). Overlap between the three set of genes showed that cluster 1 contained 59 (58%) of the known TOP-RNAs while cluster 6 contained 20 (20%, Fig. 6D). 80% of the mRNAs of cluster 1 are known to be TOP-RNAs, one hypothesis is that the similar behaviour of the cluster1 mRNAs is due to their TOP motif. We calculated the TOP local score for the 14 remaining mRNAs and found a similar distribution to cluster 1.

In conclusion, we found that translation increased for more than 70% of the mRNAs coding for ribosomal proteins, and also for genes involved in specific metabolic processes, response to growth factors or that regulate cell death.

### mTOR and eIF2alpha pathways are regulated upon glucose induction

We next sought to identify the molecular pathways driving such translation regulation in response to glucose in human beta cells. Our data unraveled a group of uORF-containing mRNAs that are translationally regulated upon glucose induction. uORF-mediated translation mechanisms involve the phosphorylation of eIF2 complex alpha subunit (eIF2α) on Ser51, which inhibits the assembly and recycling of the translational ternary complex (eIF2-GTP·Met-tRNAi), and results in a reduction in translation initiation. We therefore examined the phosphorylation status of eIF2α. Western blot analyses highlighted a dephosphorylation of eIF2α in response to glucose (Fig. 7A, P-eIF2α, and Supp. Fig. 6A for western blot quantification), indicating an increased availability of the ternary complex. Furthermore, adding the translation inhibitor cycloheximide does not prevent the activation of eIF2 through dephosphorylation, which is therefore independent of protein neosynthesis. Note that the amount of poly-A binding protein (PABP), a key post-transcriptional regulator, remains constant.

**Figure 7.**
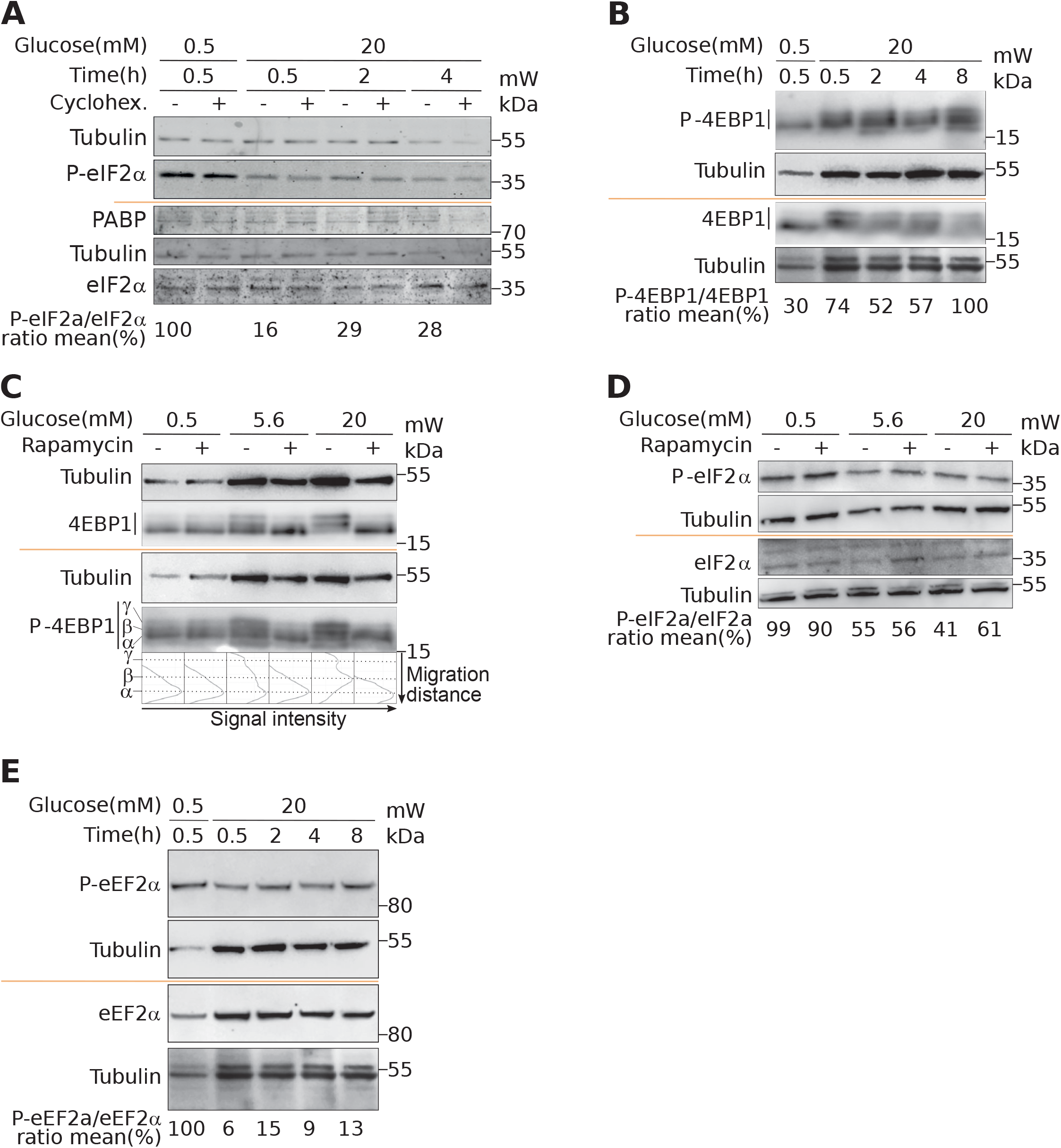
Activation of mTOR and eIF2α upon glucose induction. Western blot analysis of cells incubated with 0.5- or 20-mM glucose for the indicated time. Where indicated, cells were pre-treated for one hour with cycloheximide (CHX) to block translation, or with rapamycin to inhibit mTOR activation (see text). For 4EBP1, upper bands (β and γ) correspond to hyperphosphorylated forms. The ratio phosphorylated/total proteins was reported from quantification presented in Supp. Fig .6.

It has been described that glucose deprivation is sensed by aldolases, which cause the formation of a membrane-associated lysosomal complex, which in turn activates AMP-activated protein kinase (AMPK) (26). AMPK induces inhibition of mammalian TOR complex 1 (mTORC1) activity by phosphorylation of the tuberous sclerosis protein 2 (TSC2) tumor suppressor and the mTOR binding partner Raptor (27,28). Since mTORC1 is a well-known regulator of protein synthesis, we postulated that upon glucose increase, mTORC1 activation participates in translation upregulation. It is well established that mTORC1 regulates translation through the modulation of the phosphorylation status of 4EBP1 (29). Thus, to further validate that mTORC1 signaling is a key player in glucose stimulation, we checked the phosphorylation status of 4EBP (Fig. 7B and C). At 0.5 mM glucose, 4E-BP displayed a single band (α, see Fig. 7C) in western blot that is also visible using an anti phospho-4EBP antibody showing that 4EBP is at least partly phosphorylated. The P-4EBP/4EBP ratio increased upon glucose shift, and new 4EBP isoforms are visible (bands β and γ see profiles in Fig. 7C), corresponding to hyperphosphorylated 4EBP. We concluded that increase of glucose concentration induced strong phosphorylation of 4EBP that is known to decrease its affinity for eIF4E, and increases Cap-dependent translation. We also observed during a proteomic study dedicated to the analysis of the late response to glucose (4h after glucose increase), that RPS6 was phosphorylated rapidly after glucose addition, confirming the activation of the mTOR pathway (30). Interestingly, TOP-containing mRNAs are known to be extremely sensitive to mTOR regulation, that provides an attractive regulation mechanism to understand the translation upregulation of mRNAs of cluster 1 and 6 (see Discussion). Furthermore, using rapamycin, a potent inhibitor of mTOR, we observed that eIF2 dephosphorylation was independent of mTOR activation (Fig. 7D). Since we found the kinase eEF2K in cluster 4, we asked whether eEF2 phosphorylation was affected upon glucose shift. Indeed, we found a strong reduction of eEF2 phosphorylation (Fig. 7E, and quantification in Supp. Fig. 6D), that is known to promote its activity for translation elongation. In conclusion, from the structural features of the clusters, we focused on mTOR and eIF2α pathways that are independently regulated to modulate translation after a short glucose stimulation, and found also an activation of the elongation factor eEF2.

## DISCUSSION

We show here that a human pancreatic β-cell line responds to glucose by a specific regulation in protein synthesis, as revealed by polysome profiling. Interestingly, this increase in translation is unrelated to the abundance of mRNAs and consequently independent from a transcriptional regulation. Indeed, from our transcriptome analysis, a 30 min glucose shift had no effect on mRNA abundance, apart for 16 transcripts that are weakly affected. This result agrees with a microarray study performed in EndoC-βH1 in which only few genes had their mRNA levels affected after 8-hour of glucose stimulation (6).

This context where translational regulation is the major determinant of gene expression prompted us to perform the first translatome study of a human pancreatic β-cell line in response to glucose. Following a 30 min incubation in high-glucose media, we found changes in distribution of 402 mRNAs in different ribosomal fractions. We report that two important genes involved in the regulation of secretory granules, *CHGA* and *PTPRN* are translationally co-regulated in human pancreatic β-cells. The PTPRN translation increase was also reported in mouse models (16). PTPRN belongs to the receptor protein tyrosine phosphatase (RPTP) family and regulate basal and glucose-induced insulin secretion in the mouse MIN6 cell line by increasing, presumably through stabilization, the number of insulin-containing dense core vesicles (31). Conversely, we found that the association of CHGA with heavy polysomes decreases and consequently its translation is reduced. CHGA is a member of the granin glycoprotein family, and its main intracellular function is to sort proteins into the secretory granules. CHGA is also secreted and generate several cleaved products among which pancreastatin that have been shown to act in an autocrine and paracrine fashion by inhibiting glucose stimulated insulin secretion (32).

We observed, by clustering analysis, that the identified differentially translated mRNAs could be divided into six groups based on the different changes of their mRNA levels in the three ribosomal sequenced fractions. We thus performed mRNA features analysis to identify possible mRNA features that could explain the observed different behaviours.

It is interesting to note that transcripts from cluster 4, that are translationally repressed upon glucose shift, possess a longer 3’UTR that could contain more regulatory elements, such as miRNA binding sites. Upregulation of miRNAs have been described in the presence of high glucose (33), thus, it would be interesting to analyze the modifications in miRNA activity induced by glucose in our human cellular model.

We found mRNAs from clusters 4 and 5 better translated in low glucose where eIF2 is hyperphosphorylated and, less translated in high glucose after dephosphorylation of this initiation factor. A similar situation has been observed in mice MIN6 beta cells (see introduction, (11)): in low glucose eIF2 phosphorylation is mediated by the Integrated Stress Response (ISR) that is suppressed upon glucose increase. Indeed, we found a similar translational behaviour for ATF4, DDIT3 and c-Jun transcripts encoding proteins associated with the ISR (34): they all belongs to cluster 5. Importantly, abundance of these mRNAs decreases upon glucose increase in mouse cells but their abundance does not vary in our human beta cells.

As hyperphosphorylation of eIF2α is known to favour translation of mRNAs containing uORFs, we could expect to find uORF-containing mRNAs in cluster 4 and 5, and indeed we found mRNAs reported to contain uORFs that regulate their translation: the cyclic AMP-dependent transcription factor (ATF4), the activating transcription factor 5 (ATF5) (35), the transcriptional regulator CHOP (36) and eEF2K (37).

ATF4 was the most translationally downregulated gene identified (log2 FC -1.5 in heavy polysomes). Importantly ATF4 is known to be the master regulator of cellular metabolism in response to energetic stresses and depending on the intensity and length of the stress can either favour cell survival through upregulation of autophagy related genes and amino acid transporters or enhance expression of genes involved in apoptotic processes (38). ATF5 has been recently shown to play an important role in regulating pancreatic β-cells survival (39,40). Despite this decrease in ATF4 and ATF5, a significant transcriptomic response was not observed for 30 min (this study), or for eight hours (6) of glucose stimulation. The ATF4 protein increase is classically described to activate transcription of target genes, but this response decreases with time owing to different mechanisms that counteract ATF4 function (reviewed in (41)). The reduction in ATF4 translation would then allow the cells to return to basal levels of ATF4 without triggering a transcriptional response.

It is interesting to note that after glucose shift, translation inhibition of eEF2K may participate in translation regulation. eEF2K phosphorylates eEF2, reducing its affinity for ribosomes, resulting in inhibition of protein synthesis (42). Indeed, we observed a strong dephosphorylation of eEF2, that would promote the elongation rate of translating ribosomes, participating to the global protein synthesis increase revealed by the measure of amino-acid incorporation.

IRESs are also RNA structures conferring a translational advantage in condition where general translation is silenced. We searched in the literature if any of the translationally repressed mRNAs were reported to contain an IRES. We found that the *RRBP1* (Ribosome-binding protein 1, cluster 5) mRNA has been shown to contain an IRES in its 5’ UTR (43). RRBP1 is a membrane-bound protein found in the endoplasmic reticulum where it enhances the association of certain mRNAs (44) and play a role in ER morphology (45). Consequently, RRBP1 may participate to the reshaping of the translatome upon glucose induction.

eIF2α has been implicated in many physiological translation regulations, being a “funnel factor” where several signals converge to regulate its phosphorylation at serine 51, which results in cap-dependent protein translation repression (46), as observed for the ISR. The ISR aimed to protect cells against various cellular stresses, including viral infection, oxidative stress and ER stress. Interestingly, phosphorylated eIF2α is essential to preserve ER integrity in beta cells, and if this mechanism of protection is compromised, it would contribute to the onset of Diabetic Mellitus (47), a public health concern worldwide with an increased incidence of morbidity and mortality.

The metabolism of the beta cell is also reshaped during this early response to glucose (30 mn after glucose shift) as we found in cluster 4 transcripts coding for proteins involved in regulation of metabolic processes, cell death and response to growth factors. This translational regulation is particularly important for genes implicated in tricarboxylic acid (TCA, Krebs Cycle), acyl-CoA, and thioester metabolisms. Activation of these metabolisms, that are linked by their role in glucose consumption for energy production, is expected upon translation increase, since protein synthesis is one of the most energy costly cellular processes (48). In another study done by mass spectrometry to monitor the late response to glucose (4 hours after glucose shift), we have also observed a regulation in the amount of proteins involved in TCA metabolism and glycolysis (30).

The mRNAs that showed the strongest increase in the light polysome fraction were grouped in clusters 2 and 3. Notably these clusters showed a high GC content for both the UTR regions. The GC content in the 5’ UTR could imply a strong dependency toward helicases, and a reduced initiation activity, which could explain why these mRNAs cannot load enough ribosomes to efficiently access to heavy polysomes.

We have shown that transcripts of the two most highly translated clusters have shorter coding sequences and shorter and less complex 5’UTRs compared to the rest of the clusters. These features are characteristic of a special class of mRNAs, the TOP-mRNAs (24), which are all downstream targets of the mTOR pathway (see below). This class is defined by a 5′terminal oligopyrimidine (TOP) motif that is indeed enriched in these two highly translated clusters. Also most mRNAs from cluster 1 are known as TOP-mRNAs. The remaining 14 mRNAs of cluster 1 are most probably new TOP-RNAs, as suggested by their TOP-local score distribution. Most of the known TOP-RNAs encode proteins of the translation machinery. Accordingly, gene ontology analysis revealed that cluster 1 was enriched for categories related to structural components of the ribosomes involved in ER-related translation. Amongst the 14 new putative TOP-mRNAs that we found up-regulated in these pancreatic beta cells, we found transcripts coding for proteins acting in ubiquitin binding, cell signalling and mRNA translation. 20 mRNAs from cluster 6 were also previously reported to be TOP-RNAs (Fig. 6E). By comparing the mRNA features of the cluster 1 and 6, we noticed similar characteristics that promote translation activation, such as short and unstructured 5’UTR, but transcripts from cluster 6 have longer CDS. This feature may explain why more ribosomes are loaded on cluster 6 mRNAs (at least 4 ribosomes) for a constant initiation rate, leading to their depletion from light polysomes and an enrichment in heavy polysomes.

The TOP-RNAs have been described to be regulated by the mTOR pathway in various situations, such as changes in nutrients and other growth signals (49). Gomez and co-workers (50), studying glucose stimulation in murine MIN6 cells, concluded that translation regulation by glucose is largely independent of mTOR but mainly dependent on the availability of the ternary complex regulated by eIF2α phosphorylation status.

In our human cellular model of beta cells, both mTOR activation and eIF2 dephosphorylation participate to the increase of mRNA translational increase. It is interesting to note that the eIF2α activation is independent of the mTOR pathway since using rapamycin, a potent inhibitor of the mTOR pathway, we have still observed the dephosphorylation of eIF2. These pathways regulate mRNA translation in particular through uORF and TOP features, that are also mutually exclusive on mRNAs, meaning that these regulations occur independently.

We concluded from our results that the glucose-dependent mTOR activation have a crucial importance for the nature of the transcripts that are regulated by glucose in human cells. As a quick response to glucose increase, TOP-RNAs allow accumulation of the translation machinery to prepare the beta cells for further protein demand due to the glucose-mediated metabolism changes.

Adaptation and response to glucose of pancreatic beta cells is critical for the maintenance of normoglycemia. Its deregulation is associated with Diabetic Mellitus. Mice models, animal or cell derived models, have tremendously contributed to our understanding of human biology. All too often, however, gene expression differ markedly from human cellular models (51). Despite extensive research in rodent models of beta cells, gene expression regulation in response to glucose remained largely unexplored in human cells beta cells. Using the only human cell line available of pancreatic β-cells exhibiting glucose-inducible insulin secretion (12), our results emphasize a remarkable difference in gene expression regulation in response to glucose that occurs mainly at the transcriptional level in mouse and at a translational level in human pancreatic β-cell.

We have described the first genome-wide translatome study of a human pancreatic β-cell stimulated by glucose, highlighting that the response is translational and virtually independent from changes in mRNA abundance. Through the recognition of specific mRNA features, the swift translation activation is particularly efficient to increase translation machinery components. Finally, the combined mTOR and eIF2α activation that leads to the translatome reshaping governed by specific mRNA features allows a quick and direct cellular response targeting the translational regulation. These results constitute a call for a new paradigm of gene expression regulation to better understand β-cell glucose-mediated metabolism, encouraging biologists and clinicians, whenever possible, to complement their transcriptomic studies with analysis at the translational or proteomic level.

## MATERIALS AND METHODS

### Cell culture and treatment

EndoC-βH2 cells (12) were cultured in low-glucose (5.6 mmol/L) DMEM (Sigma-Aldrich) with 2% BSA fraction V (Roche-Diagnostics), 50 mmol/L 2-mercaptoethanol,10 mmol/L nicotinamide (Calbiochem), 5.5 mg/mL transferrin (Sigma Aldrich), 6.7 ng/mL selenite (Sigma-Aldrich), 100 units/mL penicillin, and 100 mg/mL streptomycin. Cells were seeded at a 40% confluence on plates coated with Matrigel (1%; Sigma-Aldrich), fibronectin (2 mg/mL; Sigma-Aldrich). Cells were cultured at 37°C and 5% CO_2_ in an incubator and passaged once a week when they were 90–95% confluent. For the polysome profile experiments cells were plated 4 days before treatment to reach 80-90% confluence the day of experiment. Cells were cultivated for 24 h at 0.5 mM glucose and were then treated with different concentrations of glucose to obtain media at 5.6 mM, high-glucose media at 20 mM or with low-glucose media at 0.5 mM for 30 min.

### Western blot & antibodies

Protein concentrations were quantified using a Pierce BCA Protein Assay Kit (Thermo Fisher Scientific). Proteins (20 μg) were resolved by SDS-PAGE and transferred to a membrane using an iBlot2 Gel Transfer Device (Thermo Fisher Scientific). Membranes were incubated with specific primary antibodies against: phospho-Ser52-eIF2α (SAB4300221 Sigma), eIF2α (SAB4500729 Sigma), tubulin (T9026 Sigma), phospho-4EBP1 (2855 CST), 4EBP1 (9644 CST), Phospho-eEF2 (2331 CST), eEF2 (2332, CST) and PABP1 (52). Membranes were incubated with species-specific horseradish peroxidase, or fluorescent–linked secondary antibodies (1:10,000) and visualized on a Odyssey Fc Dual-mode Imaging System Onstrument (LI-COR). Quantification was done (Supp. Fig. 6), and profiles of the Fig. 7C were obtained, using Image Studio Lite 5.2.5. These data were processed to generate graphical representation with statistics with Rstudio 1.2.1335.

### Polysome profiling

Polysome profiling was performed on three independent cell cultures both in high (20 mM glucose) or low (0.5 mM glucose) and each replicate corresponded to approximately 30 million cells. After the glucose treatment, cells were washed once in ice cold PBS containing 100 µg/ml Cycloheximide. The PBS was then removed, the lysis buffer (80 mM KCl, 10 mM Tris pH7.4, 5 mM MgCl2, 0.5% Triton X 100, 0.5% Na-Deoxycholate, 40U/ µL RNAsin, 1 mM DTT) was added directly to the plate and cells were scraped and collected. After 10 minutes incubation on ice, the lysates were centrifuge at 10,000 x g for 5 min at 4°C. 10 A254 units of lysates were layered onto a 11 ml 20–50% (wt/vol) sucrose gradient prepared in the lysis buffer without Triton X-100. The samples were ultra-centrifuged at 39,000 × g for 2.5 h at 4 °C in a SW41 rotor. The gradients were fractionated in 14 fractions of 0.9 ml using an ISCO fractionation system with concomitant measurement of A254. Polysome/monosome ratios were obtained by dividing the area of the polysomal peaks by the area of the peak for the 80S monosomes. Total lysates and fractions were supplemented with 50 µl of 3 M NH4Ac, 10 ng of Luciferase RNA (Promega), 1 µl of Glycoblue (Ambion) and 1.2 ml of ethanol. Samples were vortexed and precipitated overnight at −20 °C. The pellets were collected by centrifugation at 10,000 × g for 10 min at 4 °C, washed once in 75% ethanol and resuspended in 100 µl DEPC-treated H2O. Samples were then treated for 1 hour at 37°C with RQ1 DNase (Promega) to remove possible contamination by DNA. RNAs were isolated by acid phenol: chloroform and precipitated in 1 ml Ethanol supplemented with (supplemented with 50 µL 3M NaOAc pH 5.2, 1 µl of glycoblue). Pellets were resuspended in 20 ul DEPC-treated H2O. For sequencing equal volumes of fractions were pooled: fractions 5-6 (Monosomes), fractions 7-9 (Light polysomes) and fractions 10-13 (Heavy polysomes). Quality and quantity of pooled fraction was tested by the bioanalyzer RNA 6000 Pico kit (Agilent). Sequencing was performed by the Genom’ic platform (Institut Cochin, Paris). Libraries were prepared using TruSeq RNA Library Preparation Kit (Illumina) with rRNA depletion using Ribo-zero rRNA removal kit (Illumina) following manufacturer’s instruction. High-throughput sequencing was performed using Hiseq 2000 (Illumina) system for 75nt single-end reads.

RNAs were extracted using RNeasy mini kit (Qiagen, ref: 74104), DNAse treatment was performed with RNAse-free DNAse Set (Qiagen, ref: 79254). Equal volumes of all samples were reverse transcribed with Superscript IV reverse transcriptase (Life Technologies) for polysomes samples (Suppl. Fig 2 C and D). Reverse transcripts were obtained using RNA at 1ug/50ul with the kit High Capacity CDNA RT (Life Technologie ref: 4368814) for total RNA samples (Suppl. Fig. 1C). qPCR was done with GoTaq® qPCR Master Mix (Promega ref: A602) on ViiA 7 Real-Time PCR System (Thermo Fisher Scientific). DNA contamination was assessed omitting the RT, no significant signal was obtained. Custom primers were designed with the tool developed by Integrated DNA technologies (IDT, https://eu.idtdna.com/scitools/Applications/RealTimePCR/), and their efficiency was determined following serial dilutions of cDNA samples. Primer sequences: PTPRN, Fw (5’-3’): GTCTCCGGCTGCTCCTCT, Rv (5’-3’): GCCTGCGGTCAAATAGACA; CHGA, Fw (5’-3’); CAAACCGCAGACCAGAGG, Rv (5’-3’); TCCAGCTCTGCTTCAATGG; Cyclophilin-A primer sequences used for normalization, Fw (5’-3’): ATGGCAAATGCTGGACCCAACA, Rv (5’-3’): ACATGCTTGCCATCCAACCACT; CCNG1, Fw (5’-3’):GATATCGTGGGGTGAGGTGA, Rv (5’- 3’):TCAGTTGTTGTCAGTACCTCTATCATC; Hist1H3C, Fw (5’-3’): GCTTGCTACTAAAGCAGCCC Rv (5’-3’): AGCGCACAGATTGGTGTCTTC; Hist1H3D, Fw (5’-3’): CCATTCCAGCGTCTAGTCCG, Rv (5’-3’): TCTGAAAACGCAGATCAGTCTTGPTPRN.

### Bioinformatic analysis of Transcriptome and Polysome sequencing

Sequencing libraries were prepared from three biological replicates for both conditions (0.5 mM and 20 mM glucose). We prepared triplicates for high and low glucose conditions to produce RNA-seq libraries containing between 16-18 million reads for transcriptome and 8-14 million reads for polysomes pools. Almost 60% of reads on average were uniquely mapped to the human genome (Supp. Fig. 1D and 2A). Reads mapped to genes annotated as “protein coding genes” were kept for further analysis. Lowly expressed genes, frequently associated with high variability between replicates, were discarded. Samples from different conditions were grouped together by hierarchical clustering and PCA (Supp. Fig. 1E and 2B), ensuring reproducibility of our replicates. As described below, we thereafter proceeded with differential gene expression analysis using the limma R package (15).

#### Pre-processing

Raw fastq file obtained from the sequencer were firstly checked for their quality using FastQC and reporting with MultiQC v1.6 (53). No reads were discarded nor trimmed.

#### Mapping, read counts and TPM calculation

Quality controlled reads were then mapped to the human genome (GRCh38 from Ensembl 92) using STAR 2.6 (54) by using the parameter --quantMode GeneCounts to generate gene counts tables. STAR aligner was further instructed to generate an output (--quantMode TranscriptomeSAM) suitable as an input for RSEM (55). RSEM reports tables with transcript per million (TPM) for genes and mRNA isoforms (56). For all the rest of downstream analyses, the tables were filtered to retain only the genes which are annotated as protein coding in the Ensembl 93 annotation tables.

#### Filtering of lowly expressed genes

Gene counts table were transformed in log CPM (Counts per Million base). Genes whose CPM values were smaller than 1 at least in one sample were discarded. Then a customized R function further filtered genes whose coefficient of variation (defined as the ratio of the standard deviation over the mean) within replicates was lower than 0.75 and the mean CPM expression was higher than 4. As a first step for quality control of our datasets we performed a hierarchical clustering analysis by using the TPM tables for each sample. Clustering was performed on the Euclidean distance matrix and the Ward’s minimum variance method was used for forming clusters (option Ward.D2 in the hclust function of R).

#### Differential expression analysis and clustering

Analyses were performed using the limma (15) Bioconductor package. Differentially expressed genes (DEGs) or translated genes (DTGs) were identified by fitting linear models between all the pairs of the three polysome profile fractions applying the ebayes method to calculate p-values. Only genes with adjusted p-values for multiple testing ≤ 0.05 were selected. A separate list containing the TPM (transcripts per million reads) values was kept for downstream analysis. The average expression of each gene was calculated in each fraction (monosomes, light polysome and heavy polysomes) and condition (low or high glucose). Then for each fraction and gene the logarithmic ratio of means of high glucose over low glucose was calculated (log ratio of mean TPM expression). The generated log ratio matrix was then used in our integrative clustering approach which comprised the application of 3 clustering algorithms. Hierarchical clustering (hclust), k-means clustering (kmeans) and a model based bayesian approach clustering (mclust) were applied to the log ratio matrix of translation. Based on the silhouette measure for each clustering we evaluated that the mclust method represents better the structure of the translation data set.

#### mRNA features collection

We developed an in-house software for the retrieval and calculation of an array of sequence features for a given set of genes, The RNA features extraction tool is freely accessible in the GitHub repository (https://github.com/parisepigenetics/rna_feat_ext). The tool searches for either the most well annotated transcript for each gene in the gene list or it can also choose the most expressed transcript if a transcript abundance file is provided (e.g. the output of tools like RSEM or StringTie). We used the latter possibility using the average of all the polysomal samples. For each transcript, the tool extracts different mRNA features using the bioMart API. The mRNA features that we extracted were: length of the 5’UTR, CDS and 3’UTR and the GC content of both 5’ and 3’ UTR. The software also calculates the folding free energy for the 5’ and 3’ UTR (by using the RNAfold algorithm of the Vienna package (57), normalized by the length (MFE per bp) that is a measure of the stability and the complexity of the RNA secondary structure, an “in-house” devised TOP-mRNA local score and the Codon Adaptation Index (CAI, (58)), based on the codon usage of human genes. All statistical analyses of the features distributions along different translation behaviours were conducted by the groups based statistics R package *ggstatsplot* (https://cran.r-project.org/web/packages/ggstatsplot/index.html) using the Kruskal-Wallis H-test for comparing independent samples.

#### Enrichment analyses

We perform an array of different enrichment analyses as they are included in the clusterProfiler and DOSE R packages (23,59) including Gene Ontology annotation analysis (for all “biological processes”, “molecular functions” and “cellular component” categories), gene set enrichment analysis GSEA, KEGG pathway analysis (60) and Reactome pathway analysis (61). We visualise the results of the most significant enrichment on categories, gene sets and pathways by using typical bar/dot-plots of enrichment and a powerful graphical output of the above R package, the Gene-concept network plot (cneplot from clusterProfiler).

#### Translation ratio, translation efficiency

We define and calculate for translation ratio by using the measure of stable state mRNA from RNA-seq and a measure for mono/polysome fraction occupancy from polysome profile. We first computed the average of all the 6 polysome profile conditions (mono-, light-heavy-in high and low glucose) and the average of the two RNA-seq conditions (high and low glucose) and then we simply divide each polysome profile condition average with the respective RNA-seq average. This calculation resulted in 6 measurements of translation ratio for all genes in the three polysome fractions (monosomes, light and heavy polysomes) and the two glucose treatments (high and low). Then we computed what we call the translation efficiency of each gene in both glucose treatments (high and low) by subtracting the average of translation ratio in light plus heavy polysomes in high glucose from the same average in low glucose. These calculations allowed us to distinguish the most translated genes and those ones with the biggest shift in translation between high and low glucose. We have classified differences in translation ratios between high and low glucose to 3 groups according to the log fold-change (FC). First at +0.5 LogFC as highly translated in glucose, second with -0.25 logFC as lowly translated and a control group with +/-0.01 LogFC as control. We choose these thresholds in such a way as to generate 3 groups almost equal (UP, DOWN and control) sizes so that the comparative statistics will be more robust.

## ACKNOWLEDGEMENTS

We thank the staff of Genom’ic platform (Cochin Institute, Paris) for library preparations and sequencing, and Julia Morales (Station Biologique de Roscoff) for helpful discussions and for providing antibody samples.

## DATA AVAILABILITY

Raw sequencing data files and processes count tables for all transcripts/genes are available on Zenodo: DOI 10.5281/zenodo.4279599. The RNA features extraction tool, our inhouse software for the retrieval and calculation of an array of sequence features for a given set of genes is freely accessible in the GitHub repository: https://github.com/parisepigenetics/rna_feat_ext. The R-code for all the analyses in this work is available here: https://github.com/parisepigenetics/Translatome_Bcells_glucose.

## SUPPORTING INFORMATION FILES

Supplementary Data are available at Plos Genetics online. Supplementary_tables_Bulfoni.xls

## FUNDING DISCLOSURE

This work was supported by the Université Sorbonne Paris Cité (USPC) 2014 [appel à projets de recherche, 2014].

Funding for open access charge: [ANR-17-CE12-0010-01]

## CONFLICT OF INTEREST

Not declared

## TABLE AND FIGURES LEGENDS

**Supplementary Figure 1.**
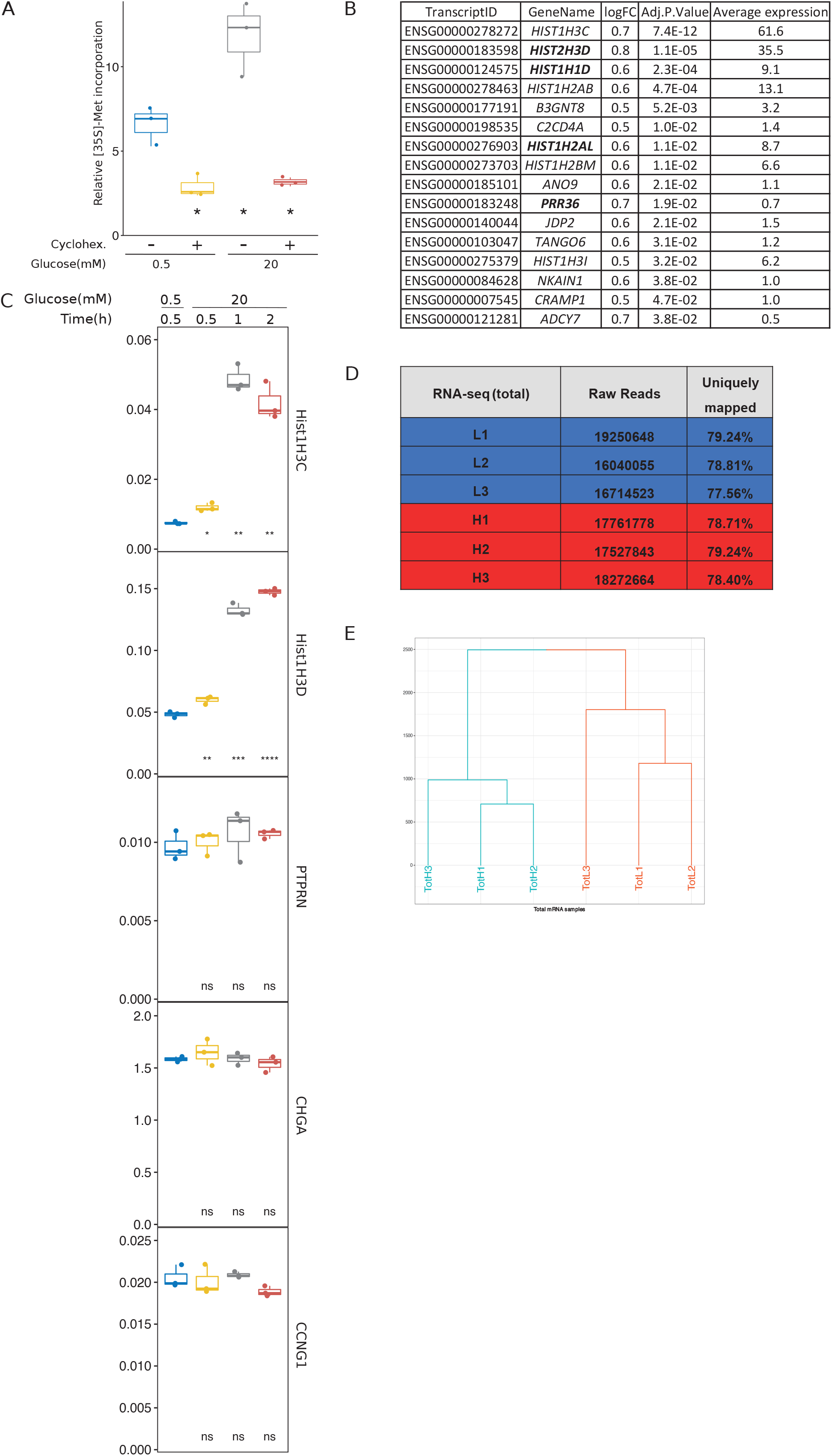
A. Translation activity increase following glucose shift observed by [35S]-Methionine incorporation. Beta-cells grown in 0.5 mM glucose were shifted to 20 mM glucose during 30 min before adding [35S]-L-methionine (ref NEG709A, Perkin Elmer, final concentration 40 microCurie/ ml) during 10 min. For each sample, [35S]-L-methionine incorporation into proteins was measured on duplicate aliquots after 10% TCA precipitation on Whatman as described in Costache et al., 2012 (PMID: 22425618). When indicated, cells pretreated with cycloheximide (Cyclohex.) that prevent protein synthesis were used as control. Numerical values of triplicates were represented as boxplots with pairwise comparison against condition 0.5mM without Cyclohex. (*, p < 0.05; Student’s t test). B. Table presenting the abundance of the 16 transcripts upregulated in high glucose condition. Transcripts also upregulated in the translatome are indicated in bold. C. RT-qPCR analysis of total RNA extracts using cyclophilin as reference gene. Delta Ct values of triplicates are represented as boxplots with pairwise comparison against condition 0.5mM glucose (Student’s t test; ns, p>0.05; *, p<0.05; **, p<0.01, ***, p< 0.001; ****, p<0.0001). D. Table summarizing the transcriptome data of total cellular extracts, after incubation of cells with low glucose (0.5 mM, L1 to L3) or high glucose (20 mM, H1 to H3). E. Hierarchical clustering of the replicates shows clear separation between low and high glucose samples.

**Supplementary Figure 2.**
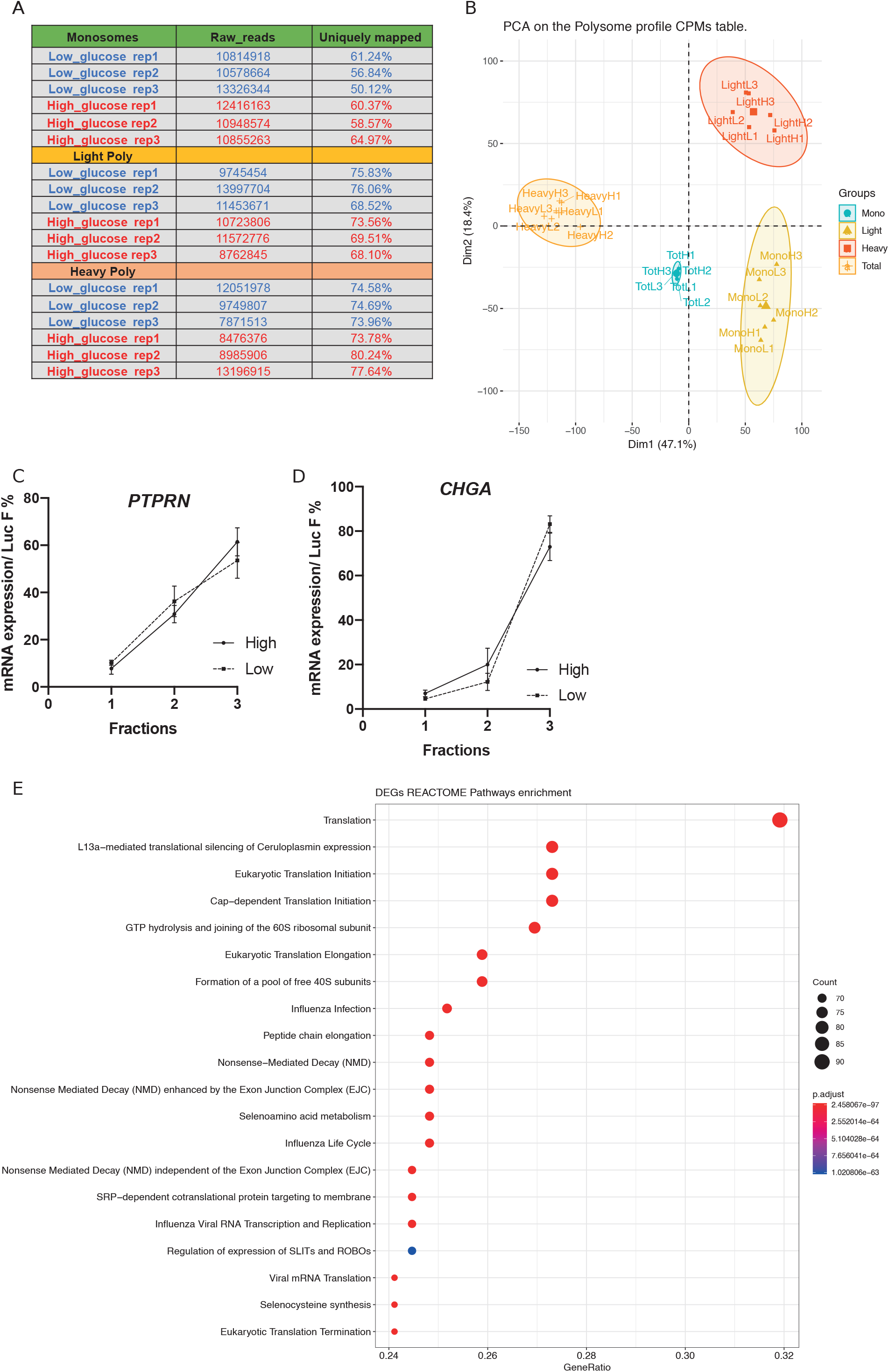
A. Table summarizing the number of raw reads and the percentage of uniquely mapped reads for each replicate in each condition. B. PCA analysis of aligned reads from each pool of fractions of the polysome profiling. C-D. RT-qPCR analysis on each of the pool of fractions. E. Dot-plot of the top 20 most significant REACTOME pathways enriched in the differentially translated genes.

**Supplementary Figure 3.**
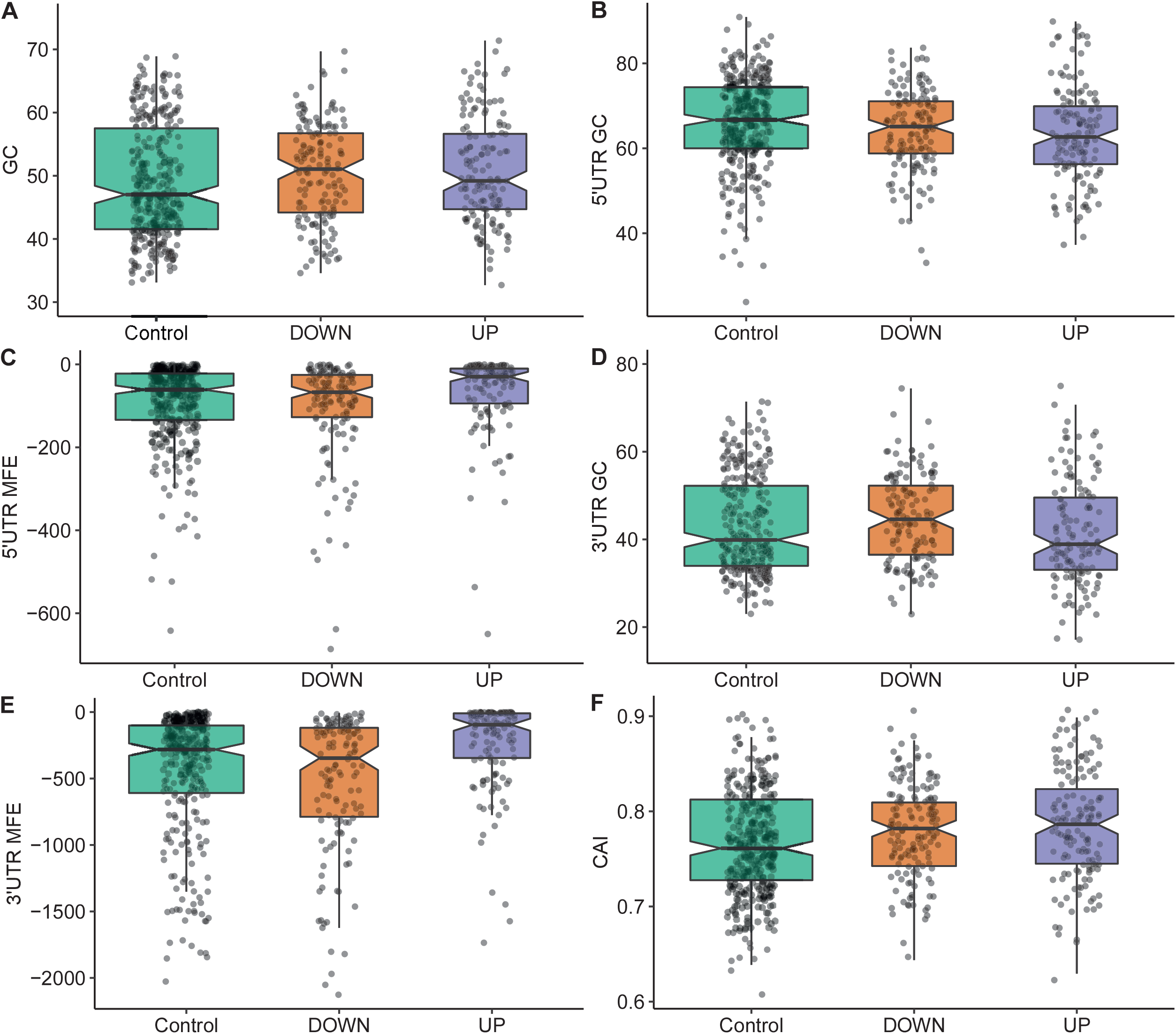
mRNA features analyses for 3 groups of mRNAs based on the translation ratio between high and low glucose (see Figure 3)).

**Supplementary Figure 4:**
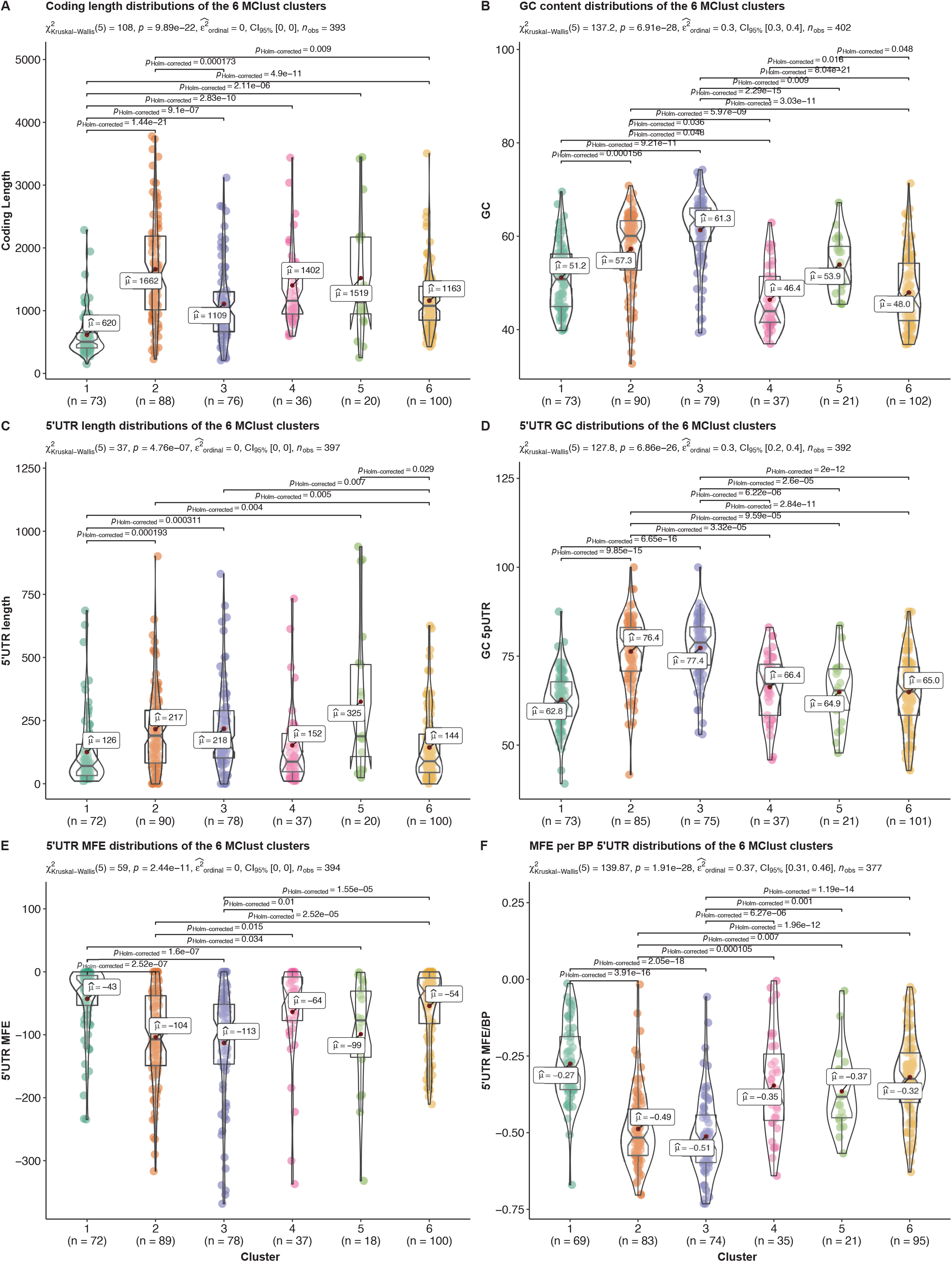
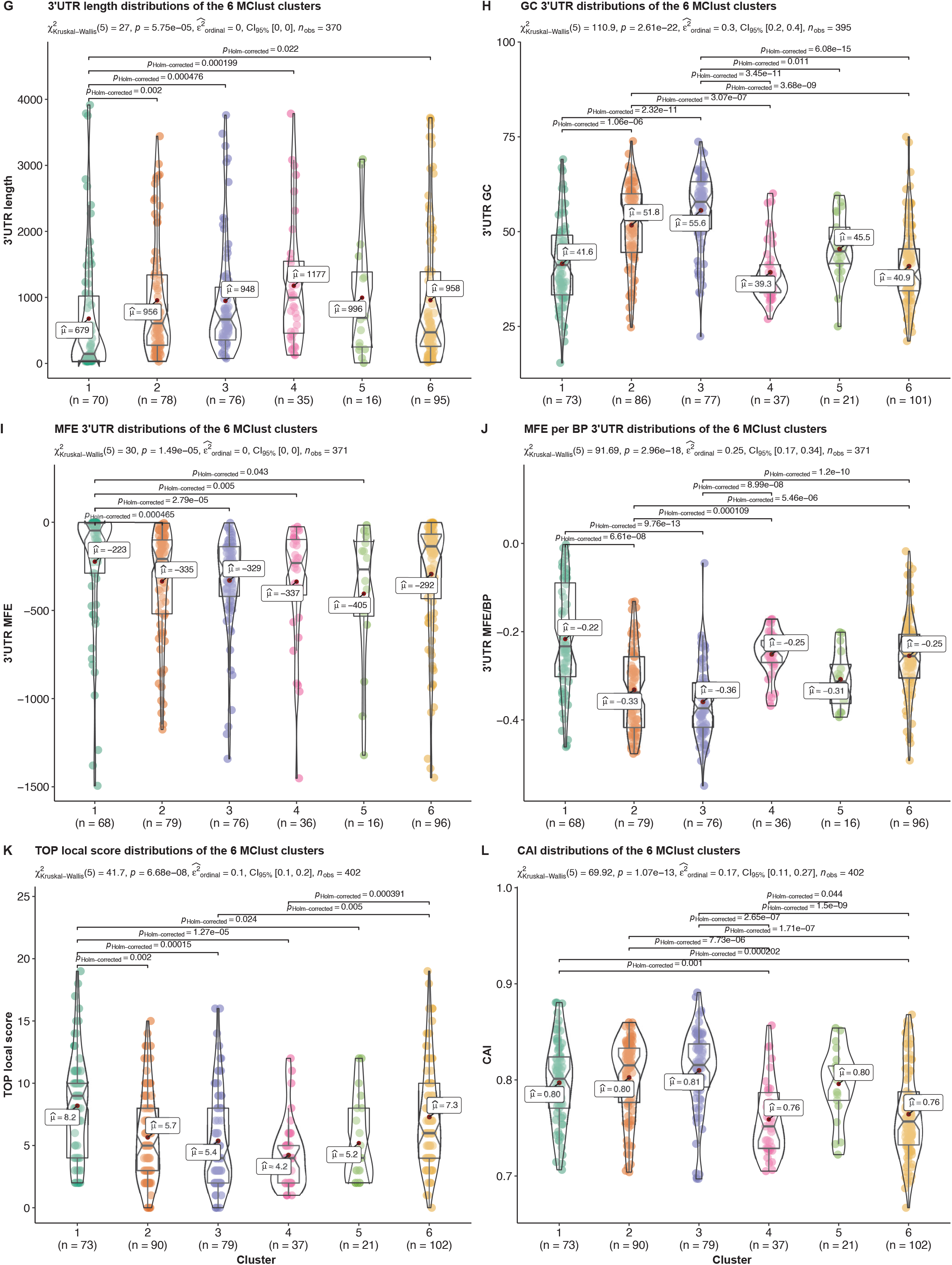
Full set of boxplots and statistical analyses for all 12 mRNA features. Each plot corresponds to an analysis of a particular feature for all the clusters of translation beha iour. he rus al allis test is used for all the group comparisons to test for median differences with the unn test for each pairwise comparison. he test statistics are shown in the subtitle and each significant pairwise comparison (corrected for multiple testing) is plotted only for significant pairwise differences

**Supplementary Figure 5.**
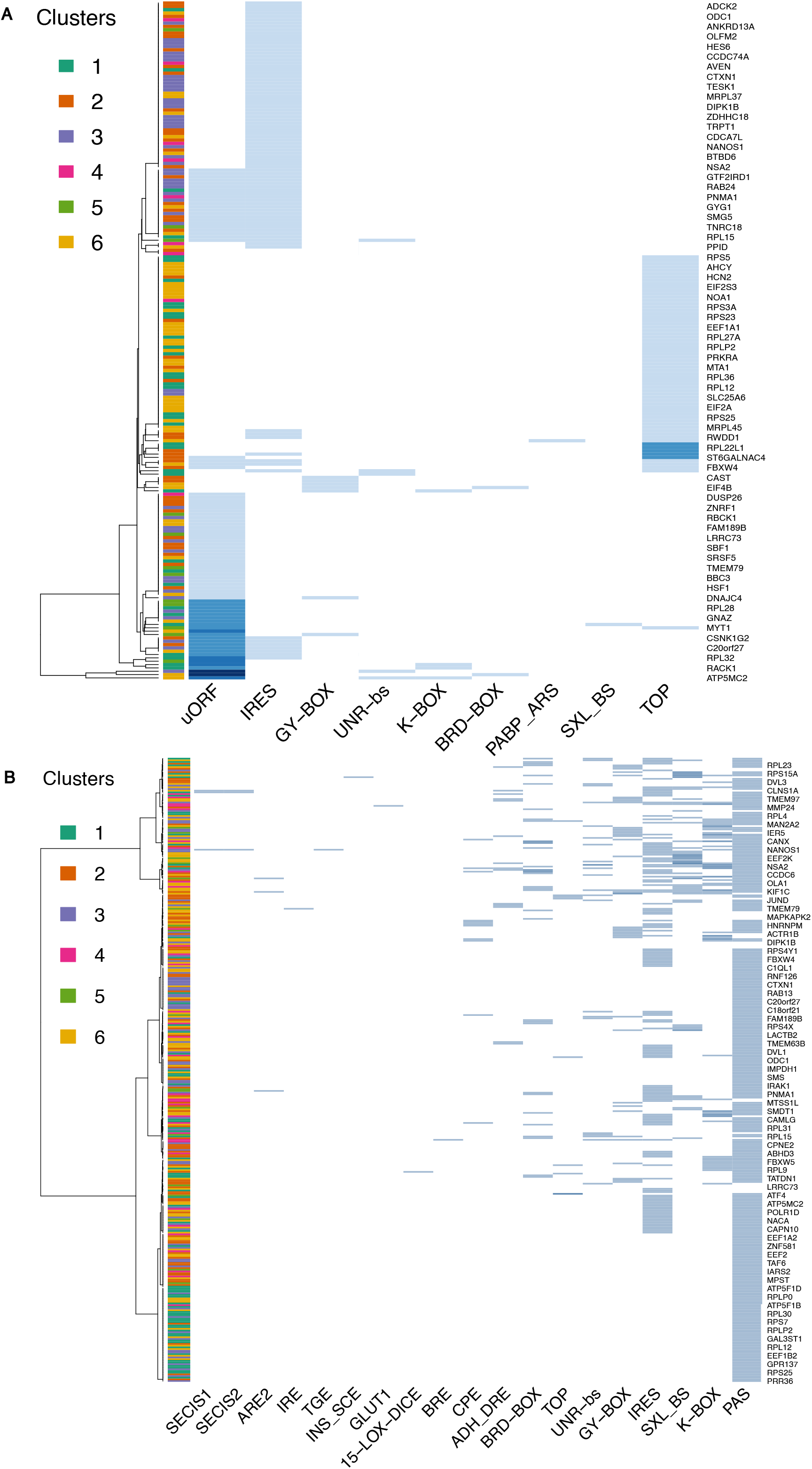
Functional motifs analysis of the 5’UTRs (A) and 3’UTRs (B) by interrogating the UTRdb. Ensembl IDs of transcripts are indicated on the right side, ordering of the transcripts was done by clustering of features.

**Supplementary Fig. 6:**
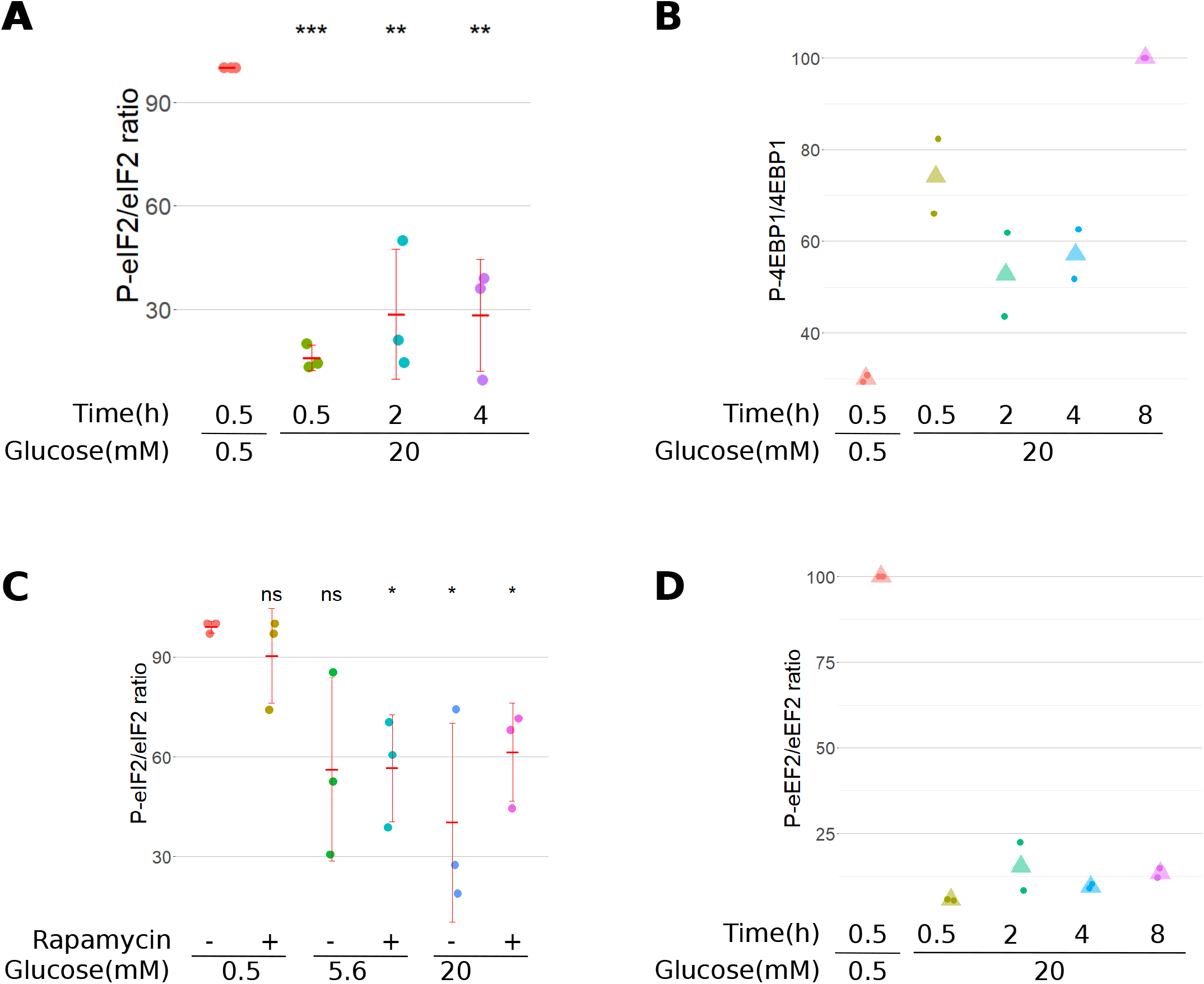
Quantification of the Western Blots of Fig. 7: A (data from Fig. 7A), C (from Fig. 7D): Numerical values of triplicates are shown along with the mean (horizontal red dash) +/- SD (vertical red line) with pairwise comparison against condition 0.5 mM (Student’s t test; *, p<0.05, **, p< 0.001, ***, p<0.0001). B (from Fig. 7B), D (from Fig. 7E): Numerical values of duplicates are shown along with the mean (triangles).

## Notes

### Competing Interest Statement

The authors have declared no competing interest.

